# Microstructural dynamics of motor learning and sleep-dependent consolidation: a diffusion imaging study

**DOI:** 10.1101/2023.05.30.542891

**Authors:** Whitney Stee, Antoine Legouhy, Michele Guerreri, Thomas Villemonteix, Hui Zhang, Philippe Peigneux

**Affiliations:** UR2NF-Neuropsychology and Functional Neuroimaging Research Unit affiliated at CRCN – Centre for Research in Cognition and Neurosciences and UNI - ULB Neuroscience Institute, Université Libre de Bruxelles (ULB), Brussels, Belgium; GIGA - Cyclotron Research Centre - In Vivo Imaging, University of Liège (ULiège), Liège, Belgium; Department of Computer Science & Centre for Medical Image Computing, University College London (UCL), London, UK; Laboratoire Psychopathologie et Processus de Changement, Paris-Lumières University, Saint-Denis, France

**Author notes:** Corresponding author: Whitney Stee Address: UR2NF - Neuropsychology and Functional Neuroimaging Research Unit, CRCN Université Libre de Bruxelles (ULB), Av. F. Roosevelt 50, CP 191, 1050 Bruxelles, Belgium.

**Keywords:** diffusion imaging, NODDI, DTI, sleep, motor memory, sequential motor learning, structural plasticity, microstructure

## Abstract

Diffusion-weighted magnetic resonance imaging (DWI) allows the observation of (micro)structural brain remodelling in cortical and subcortical regions after time-limited motor learning. Post-learning sleep consolidation should lead to long-term microstructural brain changes, but concrete evidence remains limited. Here, we used both conventional diffusion tensor imaging (DTI) and Neurite Orientation Dispersion & Density Imaging (NODDI), that estimates dendritic and axonal complexity in white and grey matter, to investigate the microstructural brain mechanisms underlying time- and sleep-dependent motor memory consolidation. Sixty-one young healthy adults underwent 2 DWI sessions, with sequential motor training in between, on day 1 followed by the experimental night of total sleep deprivation or regular sleep. After 3 recovery nights at home, they underwent 2 further DWI sessions separated by a motor retraining session. Sequential motor learning resulted in immediate modifications in structural parameters in occipitoparietal and temporal regions, as well as in subcortical regions of interest. Similar changes were observed following relearning but at a smaller magnitude. Regarding delayed consolidation effects, learning-related changes only partially persisted 3 days after initial learning, and no post-learning sleep effect was detected. Our results show rapid motor learning-related remodelling, reflecting temporary processes in learning-related neuronal brain plasticity. Post-learning sleep-related cellular changes remain to be evidenced, possibly using more sophisticated brain imaging measures and spanning more extended timescales to allow the expression of structural changes.

## 1. Introduction

Confronted with novel environmental stimulations, the brain progressively adapts both its function and structure to more efficiently meet external demands [1], [2]. In the minutes to hours following exposure to new learning material, synaptic plasticity takes place, involving early cellular determinants of synaptic strength and persistence triggered within individual neurons (e.g., changes in dendritic length, spine density or synapse formation) [3]. Additionally, modifications in glial activity can also be observed in response to learning [4]. Later on, memory consolidation takes place at the system level, where novel memories are progressively shaped and integrated into pre-existing brain networks over extended periods of time (days to weeks) including sleep [5]–[7]. For instance, rapid [8], [9] and delayed [9]–[12] changes within functional networks in response to motor skill learning have been well documented, highlighting a progressive reorganization from the beginning of the learning episode to delayed retest. As compared to the available data for functional reorganization, evidence regarding structural remodelling is less documented. Indeed, imaging studies probing experience-dependent grey (GM) or white matter (WM) structural changes (for a review, see e.g., [13]) have been limited by the relatively low sensitivity of structural magnetic resonance imaging (MRI) to detect subtle changes in the underlying brain anatomy. Therefore, only substantial changes over long and sustained training periods could be observed. For instance, the first longitudinal human study reported transient bilateral expansion in GM in the mid-temporal area (hMT/V5) and in the left posterior intraparietal sulcus after months of juggling practice [14]. GM changes in the mid-temporal area (hMT/V5) were robustly replicated in older adults using the same protocol [15]. Later longitudinal human studies demonstrated learning-induced plasticity changes in GM after a few weeks [16] or days [17] of motor skill practice. Changes in WM integrity were also shown to parallel the increased GM density in the intraparietal sulcus after repeated juggling practice [17], and correlations were found between WM integrity and GM volume in brain regions functionally engaged in motor sequence performance [18]. It is actually known that momentary adaptations in functional connectivity alter structural connections, which in turn affect functional connectivity [19]. Lastly, WM structural plasticity changes correlate with behavioural measures of improvement [20] suggesting that both GM and WM are capable of relatively rapid remodelling when acquiring novel information.

Besides a mere effect of time, post-learning sleep mechanisms contribute to memory consolidation [21] by promoting long-lasting changes in neural networks. Notwithstanding robust findings showing functional changes (using e.g., fMRI or EEG) in brain responses and behaviour after offline periods including sleep in human (for reviews, see e.g., [21], [22]), concrete evidence for sleep-dependent (micro)structural alterations in learning-related areas remains scarce. According to the synaptic homeostasis hypothesis [23], locally increased synaptic strength after learning is downscaled by slow oscillations during sleep to a baseline level, which is both energetically sustainable and beneficial for the consolidation of novel memory traces. In line with this theory, a diffusion tensor imaging (DTI) study highlighted decreased diffusivity in cortical GM following extended training, that reverted after recovery sleep [24]. At variance however, animal data showed post-learning sleep deprivation prevented the formation of branch-specific spines [25], suggesting that sleep rather promotes synapse formation and strengthening. Hence, there is no definitive evidence about the WM/GM micro-structural modifications that dynamically take place in the human brain during early, then delayed post-training periods, and their modulation by post-learning sleep. Fortunately, the last decade witnessed the speedy development of advanced MRI methods to combine brain macro- and micro-structure [26]. This enabled to dramatically shorten the timescale at which one can detect the structural remodelling that accompanies functional neuroplasticity and learning. Significant changes were thus observed over increasingly shorter learning episodes (e.g., 2 hours [20], [27] up to 1 hour [28]), using standard DTI measures, i.e., mean diffusivity (MD) and fractional anisotropy (FA) that relate to tissue density and fibre organization/directionality respectively. These rapid learning-related microstructural changes were initially mostly identified in the limbic system [20], [27], [29], but recent studies showed that such changes can also be detected in motor-learning related regions. For instance, subtle changes in MD were observed after only 45 min of motor practice in the left premotor cortex, the superior part of the cerebellum and the left middle temporal gyrus [30]. Subsequently, decreased MD was identified in the hippocampus and precuneus after only 15 minutes of motor sequence learning [31]. Although these results demonstrate the sensitivity of DTI to detect rapid and subtle changes in the underlying brain structure, there is an inherent limitation in standard DTI measures, such as MD and FA markers, that provide only aggregate information about the underlying cellular processes [32]. For example, a reduction in FA may be caused by decreased fiber density, decreased fiber coherence, or increased free water contamination. In this respect, new biophysical models have been proposed to improve specificity by disentangling the contributions from the different compartments (hindered extra-axonal and restricted intra-axonal) of the tissue and the specific geometry of each compartment [26], e.g., CHARMED - Composite Hindered And Restricted Model of Diffusion [33] or NODDI - Neurite Orientation Dispersion & Density Imaging [32]. NODDI is one of the most widely used biophysical models as it provides a better fit to the diffusion imaging data [34]. Indeed, it can model the dispersion/fanning of axonal fibres or dendrites and disentangle the microstructural effects underlying standard DTI metrics [35], [36]. More specifically, NODDI enables the voxel-wise estimation of three metrics: the neurite density index (NDI), which gauges the packing of neurites including both axons and dendrites, the orientation dispersion index (ODI), which quantifies the angular variability of the neurites, and the free water fraction (FWF).

As stated above, currently available data indicates that microstructural brain changes underlying motor sequence learning can be evidenced already in the short term, up to tens of minutes [30], [31]. There is also robust evidence that post-training sleep contributes to the consolidation of motor memories [22]. However, concrete evidence for sleep-dependent and offline structural modifications in learning-related networks in human is still lacking. Finally, there is a need to highlight the cellular processes involved in microstructural brain changes using more specific approaches than standard DTI. Therefore, we aimed at exploring using both DTI and multi-compartment diffusion imaging analysis with NODDI (1) the microstructural changes that dynamically develop in the short term (immediate learning effects at Day 1), and (2) their modulation in the long-term (at Day 5) by the availability of sleep on the first post-training night. Also, (3) we investigated the dynamics of structural reorganization that can reshape within minutes for a previously experienced material (relearning at Day 5) [3], [29]. We hypothesized the presence of (1) rapid microstructural changes following initial motor learning, (2) delayed changes modulated by the presence/absence of post-learning sleep and (3) microstructural modifications during a relearning episode at day 5, modulated by the sleep opportunity on the post-learning night. We also collected fMRI data to distinguish motor- and sequence specific brain components involved in SRTT practice.

## 2. Methods

### 2.1. Participants

Sixty-one young, healthy participants (31 females) aged 18 – 29 years (mean age ± SD = 21.31 ± 2.26) provided a written informed consent to participate in this study approved by the Liège University Hospital Ethics Committee (approval #2020/138). They were all free of any neurological/psychiatric history, had a body mass index < 28, were EEG & MRI compatible, exhibited moderate to neutral chronotype (mean score ± SD = 54.07 ± 7.84, min = 32, max = 73, Morningness–Eveningness Questionnaire [37]), and had good sleep quality (mean score ± SD = 3.33 ± 1.24, min = 0, max = 6, Pittsburgh Sleep Quality Index [38]). Musicians and computer scientists who might exhibit high-level hand motor dexterity, smokers and individuals exposed to jetlag within the past 3 months were excluded. Right- and left-handers (mean score ± SD = 4.98 ± 6.39, min = −10, max = 10, Edinburgh Inventory [39]) were both included considering the bimanual character of our motor task (see below). After being pseudo-randomly assigned to one of the 2 groups to maintain gender balance, the sleep deprivation (SD) group counted 31 participants while the regular sleep (RS) group counted 30 participants (see supplementary material Table 1 for details).

### 2.2. General Procedure

To prevent for a hormonal bias on motor performance, sleep and consolidation, women were tested during their luteal phase [40], [41]. All participants were explicitly asked to refrain from drinking caffeine or other stimulating drinks on testing days, and to maintain a regular sleep-wake schedule for the entire duration of the experiment. The regularity of the sleep–wake schedule was controlled for the entire procedure using self-reported daily sleep logs for sleep quality and duration (St. Mary’s Hospital sleep questionnaire [42]) and visual inspection of actimetric recordings (ActigraphTM wGT3X-BT, Pensacola, FL, USA).

Figure 1 illustrates the experimental design. Three days before the first testing day (Day 1), participants came to the lab for a habituation night sleeping with a 256-channels high-density EEG (hd-EEG). During the Day 1 session held around 16:30, a first baseline diffusion weighted MRI (DWI1) was acquired. Immediately after, and outside of the scanner, participants were trained for 30 blocks (approximate duration 1h) on a motor serial reaction time task (SRTT) adapted from [43], [44] (see section 2.3. for more details). Thirty minutes after the end of the SRTT session, post-training diffusion weighted MRI (DWI2) was acquired again. Participants were then informed about their assignment to one of the two possible conditions for the post-learning night, i.e., Regular Sleep (RS) or Sleep Deprivation (SD). Around 21:30, participants from the RS group were equipped with the 256-channels hd-EEG, and then slept from their regular bedtime in the sleep laboratory for the whole night (approx. 8-9 hours). SD participants spent the night awake in the laboratory for a period of 10h (maximum two participants at a time), during which they were allowed quiet activities (e.g., playing board games, read, watching non-arousing movies) under the experimenter’s supervision. Keyboard typing activities were forbidden to prevent motor interferences. Isocaloric food portions and water ad libitum were available all night. SD participants filled in hourly the Karolinska Sleepiness Scale [49] to document the evolution of their sleepiness [45], and performed every 2 hours on the 10-min version of the Psychomotor Vigilance Task (PVT [46]) to track vigilance modifications over the SD night. On Day 2 around 09:30, all SD and RS participants performed a short SRTT-retest (2 sequential blocks; approx. 2 min) and were sent back home for the 3 following nights with the instruction to keep a regular sleep-wake schedule and avoid daytime naps. In the afternoon of Day 5 (same time as Day 1 to control for circadian effect on diffusion images [47] and cognitive performance [48]), diffusion weighted MRI (DWI3) was acquired first. Then, participants were trained again for 20 blocks on the previously learned SRTT sequence (approximate duration 40 min), followed 30 min after the end of practice by diffusion weighted MRI (DWI4), and a final task-related fMRI acquisition during which they alternated motor practice on sequential and random SRTT blocks (see below). To control for between-groups or -sessions changes in behavioural alertness, participants performed the 5-min version of the PVT [49] before each SRTT session.

**Figure 1.**
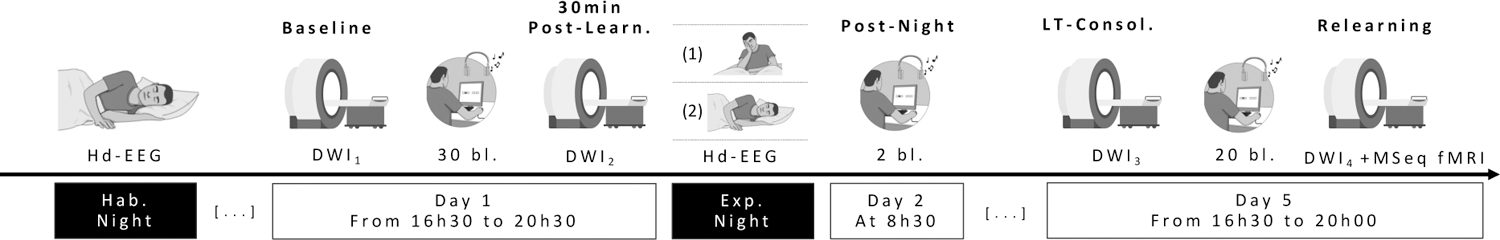
Experimental design. Three days before the first testing day, all participants come to the lab for a first habituation night under hd-EEG. On Day 1, participants undergo a first diffusion weighted imaging (DWI1) session before being trained for 1h00 on a motor sequence learning task (SRTT) in which each response key is associated with a specific auditory tone. Thirty minutes later, a second DWI (DWI2) session follows. During the subsequent night, subjects are either fully sleep deprived (1; SD) or get a regular night of sleep (2; RS) under hd-EEG. On the next morning, they all have a short behavioural retest. After 3 nights of regular sleep at home, participants come back for a 3^rd^ DWI session (DWI3). Next, they are trained again on the SRTT for 40 min followed by a last DWI session (DWI4) and then task-related fMRI during which they were asked to perform the same SRTT task in the scanner for 20 min with a pseudo-random alternation between sequential and random blocks.

### 2.3. Motor learning

#### 2.3.1. Serial Reaction Time Task

We used a 6-choice version of the SRTT (Figure 2; adapted from [43], [44]) coupled with auditory tones, running on PsychoPy3 v2020.2.10 (Nottingham, UK). Auditory coupling was established for the sake of another experiment. Participants were seated in front of a computer screen and asked to place their 6 middle fingers (index, middle and ring fingers of each hand) on 6 response keys matching the 6 squares horizontally arranged on the screen. They were given the instruction to press as fast and as accurately as possible the corresponding key every time a visual cue appeared at one of the 6 positions on the screen. Every time a key was pressed, a sound (beep tone) coupled with the key/position was emitted, and the next trial was presented after 500 msec. Ninety-six trials were presented during one block, and all blocks were separated by a short resting period (self-defined duration). The 96 trials in each block could either repeat a 12-element sequence (5-3-1-6-2-4-1-5-2-3-6-4) or be a pseudo-random succession of cues (the only restriction being that the same key is never pressed twice in a row). The learning session comprised 30 blocks (all sequential except block 26 random), the morning retest 2 sequential blocks and the relearning session 20 blocks (all sequential except blocks 3 and 16 random – also numbered as blocks 35 and 48). Random blocks were inserted to enable discrimination of respectively the sequential and motor contribution to performance improvement. Indeed, reaction time is expected to increase when there is no predictability in the succession of the stimuli (random block), indirectly demonstrated the learning of the regularity in sequential blocks.

**Figure 2.**
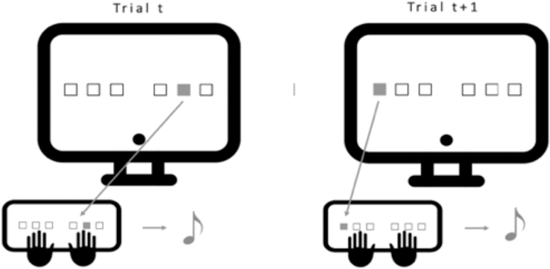
Serial Reaction Time Task (SRTT). Volunteers are seated in front of the computer screen with their 6 fingers (no pinkies and thumbs) placed on the 6 keys matching the 6 positions on the screen. Every time one of the 6 squares lights up, the participant has to press as fast and accurately as possible the corresponding key. Each time a key is pressed, an auditory tone matching this position/key is played before the apparition of the next cue (ISI = 500msec). One block is composed of 96 trials (either sequential or pseudo-random).

For the task-based fMRI session at the end of the experiment, aimed at evidencing motor learning-related networks, participants performed in the scanner the same SRTT task as described above for 30 blocks (20 sequential; 10 random) presented in a semi-random order with no more than 3 times the same block type in a row. To increase variability in the fMRI block design, half of the blocks counted 24 keypresses, the other half 36. Each block was separated from the next by a randomly determined rest duration ranging 5 to 15 s.

#### 2.3.2. Behavioural data analyses

At the behavioural level, SRTT performance was assessed for each block computing the mean reaction time (RT) and accuracy (percentage of correct triplets throughout the 96 keypresses, as humans show a natural tendency to divide behavioural sequences in chunks [50], [51] up to 3 elements [52]). Frequentist statistics were computed using JASP version 0.15 (JASP Team (2021)). Welch t-tests and Welch ANOVAs were always preferred to Student t-tests and classical One-way ANOVAs considering their increased power in case of heterogeneity of variance, that Levene’s test for equality of variances often fails to detect [53], [54]. When normality was violated, Mann–Whitney U-tests were performed. Degrees of freedom were corrected with Greenhouse–Geisser sphericity correction in case Mauchly’s sphericity test indicated violated assumption. Bonferroni correction for multiple comparison was applied when post-hoc tests were conducted. All tests are based on a two-sided significance level set at *p* < 0.05.

### 2.4. MRI Data Acquisition

MR data were acquired on a Siemens Magnetom Prisma 3T (software: Syngo MR E11) scanner. High resolution structural images were acquired for anatomical reference. Parameters for the 3D T1-weighted magnetization-prepared rapid gradient echo (MPRAGE) were acquisition time = 4 min 10 s, echo time (TE) = 2.19 ms, repetition time (TR) = 1900 ms, inversion time (TI) = 900 ms, flip angle = 9°, voxel size = 1 × 1 × 1 mm^3^, and matrix dimensions = 224 × 240 × 256 (sagittal, coronal, axial). For the 3D T2-weighted spin-echo, acquisition time was 8 min 27 s, TE = 5.66 ms, TR = 3200 ms, flip angle = 120°, voxel size = 0.7 × 0.7 × 0.7 mm^3^, and matrix dimensions = 256 × 320 × 303 (sagittal, coronal, axial). Multi-shell diffusion acquisitions were composed of 13 b = 0 and diffusion-weighted images with b-values 650, 1000 and 2000 s.mm^-2^, respective number of directions = 15, 30, 60. For distortion correction purpose, two sets of DWI acquisitions were acquired with the same settings except for the phase encoding direction (PED) that is reversed - antero-posterior (AP) and postero-anterior (PA). For the 2D axial spin-echo echo-planar imaging used for DWI, acquisition time (for one set of DWIs) was = 8 min 12 s, TE = 70.2 ms, TR = 4070 ms, flip angle = 90°, voxel size = 2 × 2 mm^2^, slice thickness = 2 mm, slice dimensions = 96 × 96 (sagittal, coronal), number of slices = 70. Lastly, for the task-based fMRI, multi-slice T2*-weighted functional images using axial slice orientation and covering the whole brain were acquired with gradient-echo echo-planar imaging (EPI), TE = 30 ms, TR = 2260 ms, flip angle = 90°, voxel size = 3 × 3 × 3 mm^3^, 25% interslice gap, number of slices = 36, matrix dimension = 72 × 72 × 36.

### 2.5. MRI Data Processing

#### 2.5.1. Anatomical processing

Raw T1-weighted images were corrected for bias field signal using BiasFieldCorrection_sqrtT1wXT2w script from https://github.com/Washington-University/HCPpipelines/tree/master, as described in the minimal preprocessing pipelines for the Human Connectome Project [55]. Segmentation was performed on the T1-weighted images using FastSurferCNN [56], an advanced deep learning model trained to replicate Freesurfer DKT’s segmentation. It segments the brain into 95 cortical and subcortical regions following the Desikan-Killiany-Tourville protocol [57], [58]. Cortical surface reconstruction was performed on T1-weighted images using FastSurfer [56] which is an extensively validated pipeline to efficiently mimic Freesurfer recon [59]–[61] by leveraging FastSurferCNN output.

#### 2.5.2. Diffusion preprocessing

The susceptibility distortion field was estimated through registration of the raw AP and PA reversed phased encoded b=0 volumes using FSL TOPUP [62]. Eddy-current distortion and head motion parameters have been estimated using FSL EDDY [63]. The reconstruction of the undistorted DWI volumes from all the raw AP and PA reversed phase encoded images was performed with the same tool through the least-squares approach from [62]. By feeding TOPUP outputs to EDDY, all the distortion and movement parameters were composed to be applied all at once, thus avoiding unnecessary resampling.

#### 2.5.3. Diffusion model fitting

The DTI model was fitted through linear least squares using FSL DTIFIT. To limit the effect of non-Gaussian diffusivity which gets stronger with high b-values [64], only the pre-preprocessed DWI volumes with b-values 0, 650 and 1000 s.mm^-2^ have been used for the fitting. The revised version [65], [66] of the original NODDI model [32] was fitted using the NODDI matlab toolbox (http://mig.cs.ucl.ac.uk/index.php?n=Tutorial.NODDImatlab). All the preprocessed DWI volumes were used for the fitting.

#### 2.5.4. Diffusion to anatomical mapping

The diffusion maps in subject native diffusion space were mapped to the high-resolution subject native anatomical space through rigid boundary-based registration [67] of the estimated b=0 image onto the T1-weighted image using FSL EPI_REG script. Diffusion metric statistics were then extracted in this native anatomical space.

#### 2.5.5. ROI-wise diffusion metrics extraction

In order to reduce the bias associated with CSF partial volume contamination when using conventional mean, we instead used the FWF estimated from NODDI to compute a tissue-weighted (“tw”) mean [68] for each ROI as summary statistic.

#### 2.5.6. Surface-wise diffusion metrics extraction

The following processing was performed using the FreeSurfer suite and outputs from the cortical surface reconstruction. Using mri_vol2surf, the diffusion metrics volumes were projected onto the mid-cortical surface, halfway through the white-grey matter border and the pial surface. Then, a smoothing kernel of FWHM 6 mm was applied along the mid-cortical surface, thus properly following the gyri and sulci circumvolutions, which usual volumetric smoothing does not allow. Surfaces of all subjects were then aligned onto a common surface template using mris_preproc.

#### 2.5.7. fMRI data preprocessing

Preprocessing was performed using the Statistical Parametric Mapping software SPM12 (Wellcome Department of Cognitive Neurology, London, UK) implemented in MATLAB R2012B (Mathworks, Sherbom, MA, USA). The four first volumes of each time series were removed to avoid residual T1 saturation effects. Individual preprocessing included realignment (2-step realignment on the first volume of the series), correction for geometric distortions caused by susceptibility-induced magnetic field inhomogeneity based on the Field Map Toolbox [69], co-registration of functional and anatomical data, spatial normalization into standard stereotactic Montreal Neurological Institute (MNI) space, and spatial smoothing using a Gaussian kernel of 6-mm full width at half maximum (FWHM).

#### 2.5.8. dMRI data analyses

For the ROI-based statistical analysis, we selected 6 bilateral subcortical ROIs based on the SRTT literature and our task-fMRI results (see below), i.e., Cerebellar Cortex [30], [31], [70], [71], Thalamus [70], [71], Hippocampus [31], [72], Caudate, Putamen, and Pallidum [70], [71]. We performed multivariate analysis of variances (MANOVA) separately on DTI (twMD, twFA) and NODDI (twNDI, twODI, FWF) parameters using SPSS version 28.0.0.0. Significance level was set at 0.008 to correct for multiple comparisons (0.05/6 ROIs). Post-hoc univariate ANOVAs were performed when necessary.

The surface-based statistical analysis was also conducted using the FreeSurfer suite. For each chosen contrast, a general linear model (GLM) was fitted on the mris_preproc outputs using mri_glmfit (different onset, different slope), and two-tailed significance for t-statistic was computed for the estimated parameters at each vertex. To account for multiple comparisons, a cluster-wise correction based on permutations [73] was performed using mri_glmfit-sim. We set 1000 permutations, a vertex-wise cluster-forming p-value (p) threshold at p < 0.001, and a cluster-wise p-value (CWP) threshold at CWP < 0.05. We did not include FA in the cortical ribbon analysis as it is not suited for GM exploration.

#### 2.5.9. fMRI data analysis

in a first-level individual analysis, a fixed-effects (FFX) model was applied to each subject’s task-based functional data, testing effects of interest by linear contrasts (difference between sequential and random blocks) convolved with a canonical hemodynamic response function and generating statistical parametric maps. Cut-off period for high-pass filtering was set at 256 s as our blocks were interleaved with 5–15 s breaks. Individual summary statistic images were then spatially smoothed (6 mm FWHM Gaussian kernel). Next, individual statistics images were introduced in a second-level analysis to evaluate differences in brain response between the SD and RS groups corresponding to a random effects (RFX) model. The resulting set of voxel values for each contrast constituted a statistical t-map (SPM(T)). Statistical inferences were obtained after correction for multiple comparisons at the voxel level (Family Wise Error (FWE) correction p < 0.05) in the whole brain. Labels were obtained using the MNI152 atlas.

## 3. Results

### 3.1. Demographic Data

Welch ANOVAs performed separately on age, laterality, sleep quality and chronotype (see supplementary Table 1) with between-subject factor *Sleep* (SD vs. RS) did not reveal any significant differences between the SD and RS groups (all *p*s > 0.388).

### 3.2. Behavioural Data

For motor learning at Day 1, mean RT progressively decreased with task practice (*p* < 0.001; see Fig. 3). A rebound in RT at pseudo-random block 26 was observed (*p* < 0.001), indicating that participants learned the sequence and started anticipating the upcoming position in sequential blocks. Thus, performance improvement was not merely due to motor practice. As expected, no group or interaction effect was found (all *ps* > 0.414) as the experimental manipulation did not happen yet at this stage. Also, accuracy remained stable for both groups over the entire learning session (all *ps* > 0.052).

**Figure 3.**
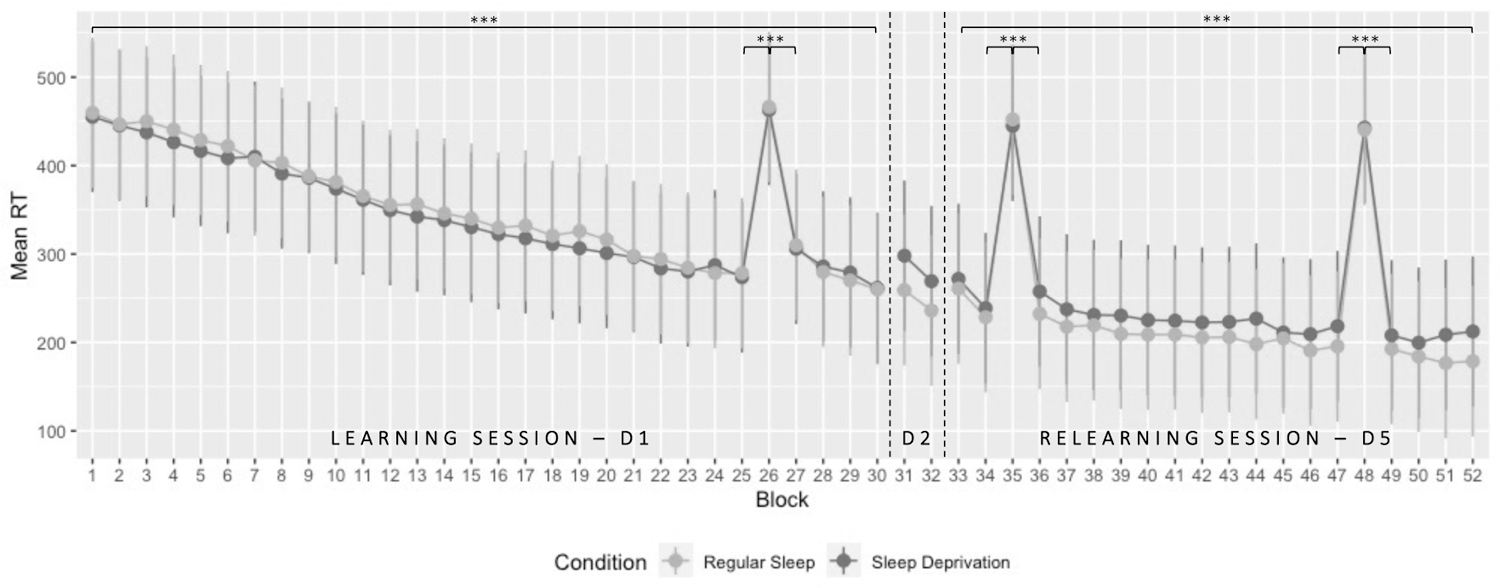
Performance evolution (speed) in the SRTT over the entire protocol. Mean RT (msec) ± standard deviation plotted for all blocks executed over the 3 different testing days. On day 1, participants performed 30 blocks (D1; block 26 pseudo-random). After the experimental night taking place between D1 & D2, performance was assessed in the morning for two blocks (D2). After three recovery nights at home, volunteers performed the task again for 20 blocks (D5; block 35 and 48 pseudo-random). *** p < 0.001.

For morning retest at Day 2, no main *Day* (*p* = 0.696) or *Group* effect (*p* = 0.354) was found when comparing the mean RT of the 2 last blocks of the learning session (LS D1; blocks 29:30) and the 2 blocks performed during retest after the experimental night (RE D2; blocks 31:32). The *Day*Sleep* interaction was significant (*p* = 0.007), but post-hoc analyses did not reveal any significant comparison after Bonferroni correction (all *p*s > 0.173). Concerning accuracy, the analysis revealed a lower accuracy at the beginning of the retest at D2 compared to the end of the learning session at Day 1 (*p* < 0.001).

For delayed motor memory consolidation at Day 5, the analysis disclosed a significant decrease (*p* < 0.001) in mean RT between the 2 last blocks of the learning session (LS D1; blocks 29:30) and the 2 first blocks of the relearning session (RL D5; blocks 33:34). Both groups exhibited a similar decrease over time (all *ps* > 0.575 for group-related effects). Also, no significant effect was found concerning changes in accuracy (all *p*s > 0.343).

Looking at motor sequence relearning at Day 5, mean RT continued to decrease over the sequential blocks (*p* < 0.001; see Fig. 3). No *Group* (*p* = 0.380) or interaction (*p* = 0.356) effect was found, suggesting that post-learning sleep availability did not impact the behavioural time course for the practice on a previously learned material. As expected, a significant RT rebound was observed at both pseudo-random blocks 35 and 48, compared to the surrounding sequential blocks (all *p*s < 0.001). Accuracy measures remained stable all over the relearning session (all *p*’s > 0.132).

We computed a complementary analysis comparing sequential-specific improvement at the end of learning and at the beginning of relearning (respectively computed as the difference in RT between the mean of the 2 sequential blocks surrounding random block 26 or 35 minus the random block, then divided by the random block and multiplied by 100) with between-subject factor *Sleep* (RS vs. SD). The analysis revealed a significant difference (*p* < 0.001) with a significantly larger difference between random and sequential bloc at the beginning of RL (−46,653%) when compared to LS (−37.219%), confirming the previous findings indicating effective delayed memory consolidation. Here again, no *Group* effect was found (all *ps* > 0.067).

Finally, all participants reported subjectively noticing the sequential nature of the task by the end of the procedure. For more detailed statistics, see section 1 in supplementary material.

### 3.3. Diffusion Weighted Imaging (DWI) Data

#### 3.3.1. Learning-related short-term structural changes (Day 1; DWI1 vs. DWI2)

##### 3.3.1.1. Cortical Ribbon

Looking at DTI parameters, the surface-based statistical analysis disclosed extended clusters in the whole sample following learning with decreased MD bilaterally in the inferior parietal and paracentral gyri, the precuneus, the insula, the precentral, lingual, superior parietal, lateral occipital, superior frontal, postcentral, supramarginal, middle-, superior- and inferior temporal gyri, the rostral anterior cingulate and fusiform gyri, the cuneus, and the transverse temporal gyrus. Smaller clusters were found in the banks of the superior temporal sulcus, the rostral middle frontal, isthmus cingulate and parahippocampal gyri as well as in the left caudal anterior cingulate and caudal middle frontal gyri, posterior cingulate and lateral orbitofrontal gyri, and right pars opercularis, pericalcarine gyrus and pars orbitalis (Fig. 4; detailed results and cluster size in supplementary material Table 2).

**Figure 4.**
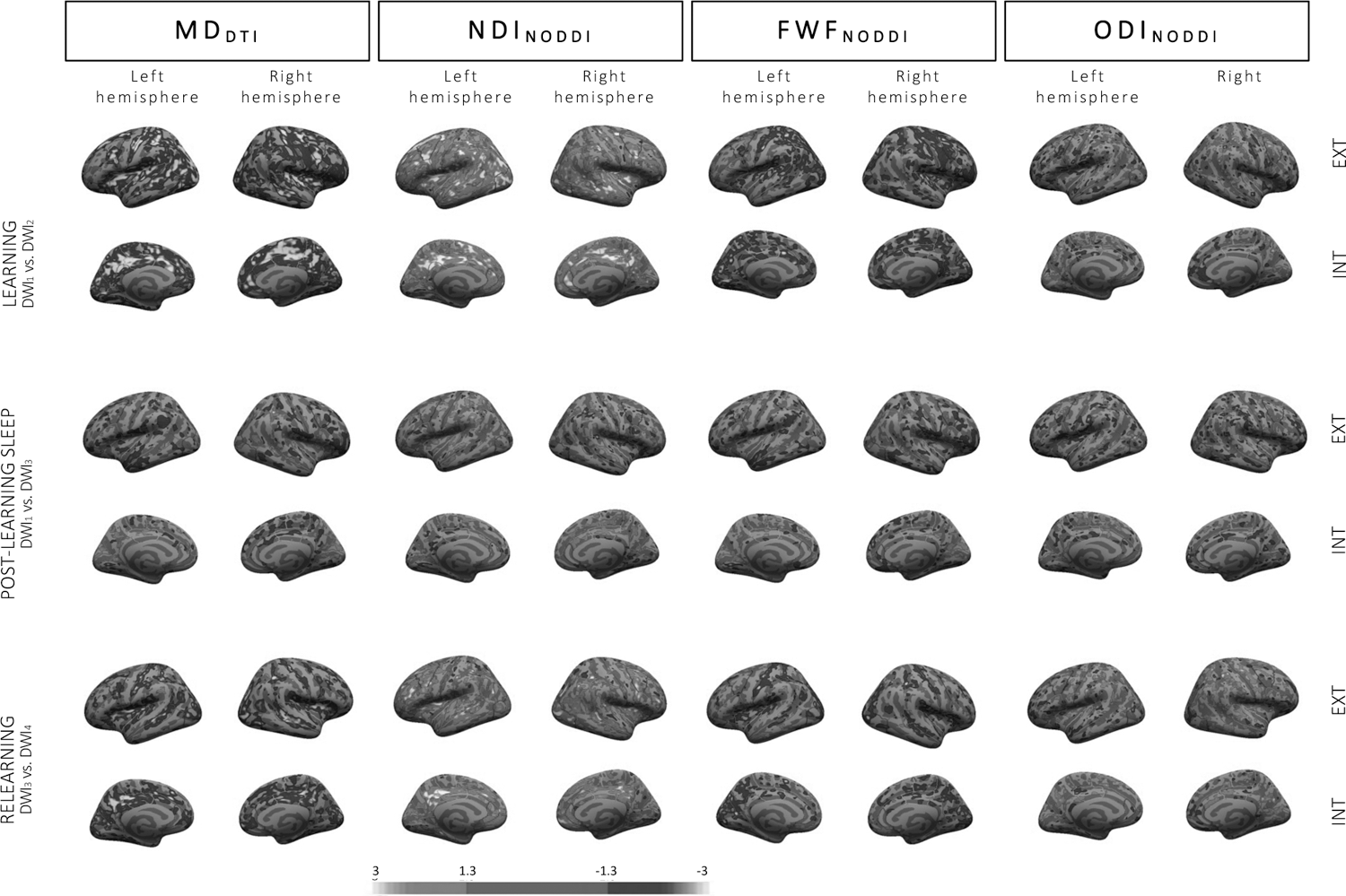
Changes in DTI and NODDI parameters in the cortical ribbon. Top row: learning-related changes in MD, NDI, FWF and ODI at Day 1 (DWI2-DWI1). Middle row: post-learning sleep-related changes (first scan at Day 5 vs. initial baseline scan; DWI3-DWI1). Bottom row: relearning-related changes at Day 5 (DWI4-DWI3). Colour-coded images depict Z-scores representation at threshold p < 0.05 uncorrected.

The surface-based analysis conducted on NODDI parameters evidenced increased NDI after learning in regions almost perfectly overlapping with regions exhibiting MD changes described above. Bilaterally, NDI increased in the lateral occipital, posterior cingulate, caudate middle frontal, inferior parietal, and lingual gyri, the precuneus, the supramarginal, superior frontal, precentral, rostral middle frontal, superior temporal, insular, postcentral, middle temporal, fusiform, and rostral anterior cingulate gyri, the pars opercularis, the lateral orbitofrontal gyrus, the pars orbitalis, the caudal anterior cingulate, medial orbitofrontal, and superior parietal gyri, the pars triangularis, and the transverse temporal gyrus. NDI clusters were also significant in the left banks of the superior temporal sulcus and cuneus and the right parahippocampal, inferior temporal, paracentral and pericalcarine gyri. FWF decrease was also evidenced in most areas exhibiting MD decrease, such as the bilateral lateral occipital, lingual, precentral, superior parietal and -frontal gyri, the cuneus, postcentral gyrus, the insula, fusiform and superior temporal gyrus, the precuneus, the caudal anterior cingulate and paracentral gyri, the left inferior parietal, pericalcarine, supramarginal, isthmus cingulate and rostral middle frontal gyri, and the right caudal middle frontal, transverse temporal, medial orbitofrontal, middle temporal, and posterior cingulate gyri. Finally, we also found ODI changes (both increase and decrease depending on the region), with small clusters in the bilateral postcentral and caudal middle frontal gyri, the precuneus, and the superior temporal and frontal gyri, left posterior cingulate, precentral, rostral middle frontal and lateral occipital gyri clusters, and right lateral orbitofrontal, inferior parietal, fusiform, supramarginal and middle temporal gyri (see supplementary material Table 2).

As expected, no between-group differences were evidenced at this stage of the procedure for any of the 4 tested metrics. Finally, we computed the correlations between DWI metrics and motor performance improvement during learning (computed as the difference in RT between the mean of the 2 last blocks (29-30) minus the mean of the 2 first blocks (1-2) divided by the mean of the 2 first blocks (1-2) multiplied by 100), as well as correlations with sequential-related amelioration (computed as the difference in RT between the mean of the 2 sequential blocks surrounding random block 26 (thus, 25 and 27) minus random block 26 divided by random block 26 multiplied by 100). No significant correlation emerged from these analyses.

##### 3.3.1.2. Subcortical ROIs

MANOVAs with within-subject factor *Learning* (Pre vs. Post) were conducted on the DTI parameters for each ROI (see Fig. 5). Significant changes were found in both sides of the cerebellar cortex (*p* < 0.001) and the hippocampus (*p* < 0.001), the right thalamus (*p* = 0.002) and caudate (*p* < 0.001), and the left putamen (*p* = 0.001). Post hoc analyses revealed a strong MD decrease in all those regions (all *p*s < 0.001), suggesting increased tissue density after learning (as compared to baseline). Also, there was increased FA in the left (*p* = 0.034) and right cerebellar cortex (*p* = 0.001), the right caudate (*p* = 0.005), and the right hippocampus (*p* = 0.035), reflecting increased directionality of diffusion inside of these regions. Additional analyses were performed to verify the absence of a *Sleep* (RS vs. SD) or interaction effect. As expected, no difference existed between groups prior the experimental night (main *Sleep* effects, all *p*s > 0.016; *Learning*Sleep* interaction, all *p*s > 0.021). Lastly, correlations between DTI metrics and motor performance were computed in ROIs exhibiting significant changes over learning. No correlation was significant for MD (all *p*s > 0.078) or FA (all *p*s > 0.302) parameters. Likewise, correlations targeting sequential-related amelioration did not disclose significant effects for MD (all *p*s > 0.226) or FA (all *p*s > 0.285).

**Figure 5.**
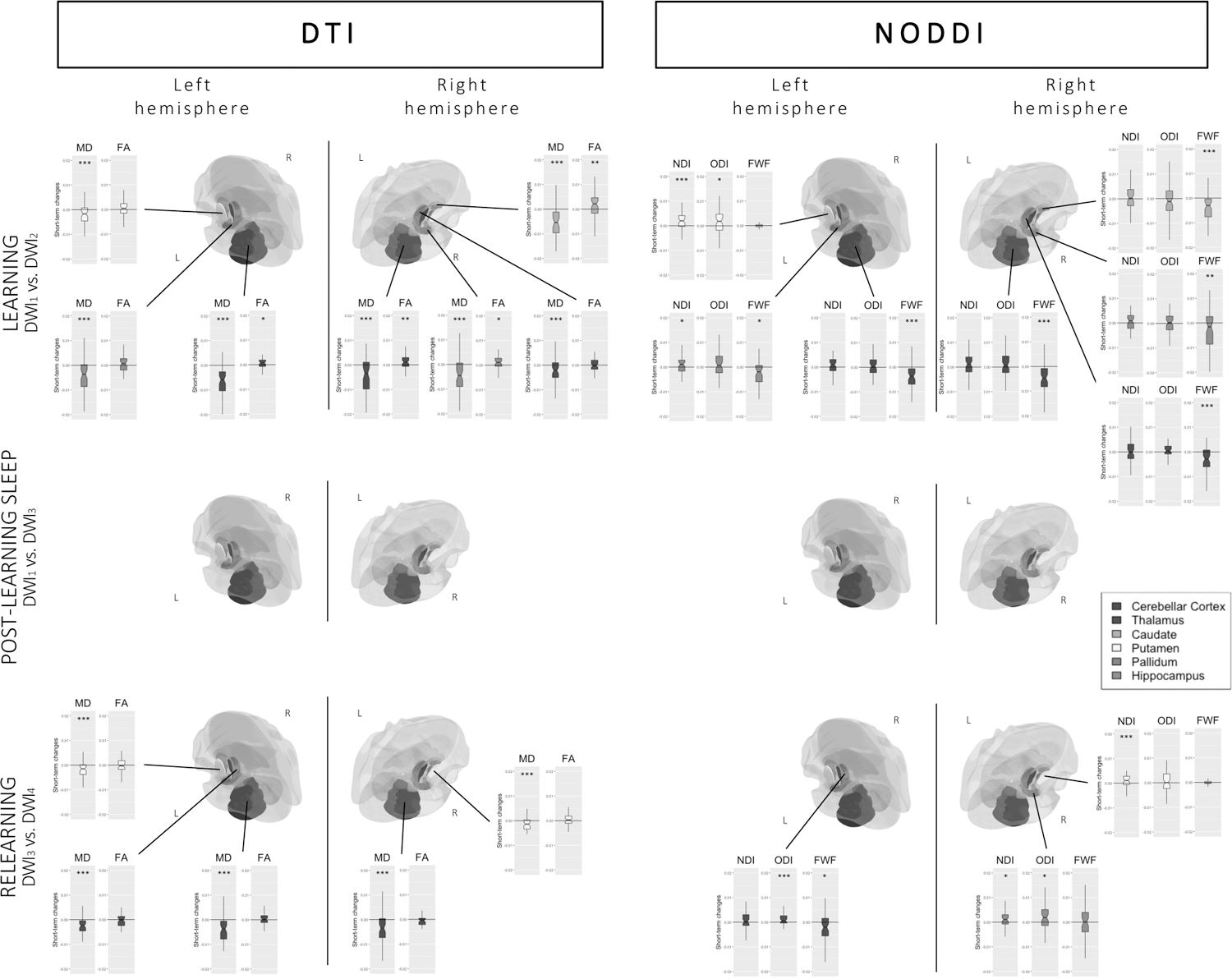
Changes in DTI and NODDI parameters in ROIs. Boxplots representing [top row] the difference between post- and pre-learning (learning effect; DWI2-DWI1), plotted for all regions in which MANOVA effects were significant, [middle row] lack of statistical differences or sleep-related interaction between baseline scan and the first scan at Day 5 (DWI3-DWI1), and [bottom row] the difference between post- and pre-relearning for all regions in which MANOVA effects were significant (DWI4-DWI3). * p < 0.05, ** p < 0.01, *** p < 0.001 for univariate post-hoc tests performed on separate metrics.

Similar MANOVAs conducted on NODDI parameters highlighted changes in similar ROIs than with DTI, i.e., both sides of the cerebellar cortex (*p* < 0.001), left (*p* < 0.001) and right hippocampus (*p* = 0.003), right thalamus (*p* = 0.004) and caudate (*p* < 0.001), and left putamen (*p* = 0.001). Post-hoc analyses disclosed decreased FWF in both sides of the cerebellar cortex (*p* < 0.001), left (*p* = 0.021) and right (*p* = 0.006) hippocampus, and right thalamus (*p* < 0.001) and caudate (*p* < 0.001), confirming the MD changes observed using DTI and suggesting increased tissue percentage. Increased NDI in the left putamen (*p* < 0.001) and left hippocampus (*p* = 0.038) suggests increased number of neurites whereas increased ODI in the left putamen (*p* = 0.048) suggests neurite reorganization and remodelling. Here again, no effect of sleep was found (main *Sleep* effects, all *p*s > 0.030; *Learning*Sleep* interaction, all *p*s > 0.034) when including *Sleep* in the ANOVA (Fig. 5). Correlations with NODDI measures in ROIs exhibiting significant changes over learning were non-significant, either with motor improvement (all *p*s > 0.254) or sequential amelioration (all *p*s > 0.157).

#### 3.3.2. Delayed structural changes and sleep-related effects in Pre-learning vs. Pre-relearning (DWI1 vs. DWI3 x RS vs. SD)

##### 3.3.2.1. Cortical Ribbon

For DTI parameters, session-related main effects were observed in the cortical ribbon with decreased MD in the left lingual and superior temporal gyri, as well as an increased MD in the right pars triangularis, the superior parietal gyrus and the banks of the superior temporal sulcus (see Fig. 4; detailed results and cluster size in supplementary material Table 3).

Looking at NODDI parameters, we found session-related NDI changes (mostly increases; see supplementary material Table 3 for detailed directionality per region) in bilateral superior temporal and lateral orbitofrontal gyri, supramarginal, rostral middle frontal and postcentral clusters, and right superior parietal and lingual gyri, pars orbitalis, fusiform and paracentral gyri, precuneus, and pars opercularis. For FWF changes, clusters mostly showing decreases were found in the bilateral fusiform gyrus, the left lingual, superior temporal and caudal middle frontal gyri and the right superior frontal, lateral occipital superior parietal, precentral and supramarginal gyri. Lastly, ODI changes were found in clusters bilaterally in the superior frontal, postcentral and inferior parietal gyri, in the left lateral occipital, lateral orbitofrontal, fusiform, precentral and supramarginal gyri and in the right rostral middle frontal and parahippocampal gyri, precuneus and inferior temporal gyrus. Sleep-related effects were non-significant, as well as correlations between behavioural performance and structural changes.

##### 3.3.2.2. Subcortical ROIs

The MANOVAs computed on DTI parameters with within-subject factor *Day* (D1 vs. D5) and between-subject factor *Sleep* (RS vs. SD) did not reveal any significant main effect of *Day* (all *p*s > 0.061), *Sleep* (all *p*s > 0.014), or *Day*Sleep* interaction (all *p*s > 0.058) for any of our ROIs. Likewise, the MANOVAs computed on NODDI parameters did not disclose main effects of *Day* (all *p*s > 0.015) or *Sleep* (all *p*s > 0.034), nor a *Day*Sleep* interaction effect (all *p*s > 0.099) in all ROIs. It suggests that learning-related changes observed at D1 in subcortical ROIs were not maintained at the beginning of D5, nor modulated by the presence of the post-learning sleep episode. Consequently, brain-behaviour correlations were not computed.

#### 3.3.3. Post-relearning structural changes and sleep-related effects (Day 5; DWI3 vs. DWI4 X RS vs. SD)

##### 3.3.3.1. Cortical Ribbon

The surface analysis comparing the post-(DWI_4_) vs. pre (DWI_3_) relearning scans did not highlight sleep-related effects.

For DTI parameters, there was a main relearning effect with MD decreases in bilateral precuneus, banks of the superior temporal sulcus, lateral occipital gyrus, insula, superior and inferior parietal gyri, cuneus, posterior cingulate, post central, superior frontal, supramarginal, lingual, caudal middle frontal, lateral orbitofrontal, precentral, superior temporal, fusiform, rostral middle frontal, and isthmus cingulate gyri. Decreased MD was also observed in the left pericalcarine and transverse temporal gyri, and in the right medial orbitofrontal, middle temporal, paracentral and parahippocampal gyri (see Fig. 4; detailed results and cluster size in supplementary material Table 4).

For NODDI parameters, increased NDI was found bilaterally in precuneus, lateral occipital, caudal middle frontal, rostral middle frontal, superior frontal and -parietal, insular, superior temporal, lateral orbitofrontal, and posterior cingulate gyri, the banks of the superior temporal sulcus, the precentral, inferior parietal, and middle temporal gyri, the pars opercularis and paracentral gyrus. NDI also increased in the left parahippocampal, postcentral, supra marginal, entorhinal, and inferior temporal gyri, and in the right medial orbitofrontal and fusiform gyri, the cuneus and isthmus cingulate gyrus. Conversely, FWF decreased bilaterally in the cuneus, lateral occipital gyrus, precuneus, postcentral, superior parietal, precentral, superior frontal, and insular gyri, in the left pericalcarine, lingual, inferior parietal rostral middle frontal, medial- and lateral orbitofrontal, and posterior cingulate gyri, and in the right -middle temporal paracentral, supramarginal, and superior temporal gyri, the banks of the superior temporal sulcus and caudal anterior cingulate gyrus. Finally, ODI changes were evidenced in the bilateral precuneus, postcentral, lateral occipital and superior frontal gyri, the left entorhinal gyrus and the right precentral, inferior, and superior parietal gyri, cuneus, insula, paracentral, rostral middle frontal, and middle temporal gyri.

Correlations between diffusion parameters and performance change over the relearning episode (computed as the difference in RT between the mean of the 2 last blocks (51-52) minus the mean of the 2 first blocks (33-34) divided by the mean of the 2 first blocks (33-34) multiplied by 100) and sequential-related improvement (computed as the difference in RT between the mean of the 2 sequential blocks surrounding random block 35 (thus, 34 and 36) minus random block 35 divided by random block 35 multiplied by 100) were non-significant.

##### 3.3.3.2. Subcortical ROIs

The MANOVA computed on DTI parameters with within-subject factor *ReLearning* (Pre vs. Post) and between factor *Sleep* (SD vs. RS) highlighted a significant main *ReLearning* effect bilaterally in the cerebellar cortex (*p* < 0.001), in the left (*p* = 0.002) and right putamen (*p* < 0.001), and in the left thalamus (*p* < 0.001). Post-hoc analyses showed that these changes were driven by decreased MD (all *p*s < 0.001) suggesting tissue densification. No changes in FA were observed (all *p*s > 0.052). There was no main *Sleep* (all *p*s > 0.022) or *ReLearning*Sleep* interaction (all *p*s > 0.020) effect, suggesting that post-learning sleep did not modulate the relearning of previously studied material. Correlations between MD and motor changes were not significant (all *p*s > 0.560), as well as with sequence learning (all *p*s > 0.268).

The MANOVA computed on NODDI parameters disclosed a main *ReLearning* effect with changes in the left thalamus (*p* < 0.001), right putamen (*p* = 0.003) and right hippocampus (*p* = 0.007). Post-hoc tests revealed decreased FWF in the left thalamus only (*p* = 0.018), increased NDI in the right putamen (*p* < 0.001) and hippocampus (*p* = 0.018), and increased ODI in the left thalamus (*p* < 0.001) and right hippocampus (*p* = 0.020). Again, there were no main *Sleep* (all *p*s > 0.040) or *ReLearning*Sleep* interaction (all *p*s > 0.063) effects. Structural changes observed in the regions mentioned above did not correlate with motor changes (all *p*s > 0.564) nor sequence learning (all *p*s > 0.141).

### 3.4. Task-fMRI Data

A two-sample t-test conducted at the random effect (RFX) level did not disclose between-group (SD vs. RS) difference in BOLD responses to sequential vs random blocks (all p_FWEcorr_ > 0.626).

A one sample t-test comparing BOLD responses to sequential vs. random blocks (over both groups) disclosed higher activity in the right (*p_FWEcorr_* = 0.001) and left precuneus (*p_FWEcorr_* = 0.001), the left hippocampus (*p_FWEcorr_* = 0.017), the left (*p_FWEcorr_* = 0.041) and right (*p_FWEcorr_* = 0.050) caudate for sequential than random blocs. Conversely, higher activity for random than sequential blocks was found in the right superior (*p_FWEcorr_* = 0.001), and middle temporal gyrus (*p_FWEcorr_* = 0.001), the right (*p_FWEcorr_* < 0.011) and left middle frontal gyrus (*p_FWEcorr_* = 0.005), and the left post central gyrus (*p_FWEcorr_* = 0.023).

### 3.5. Additional (control) analyses

For detailed analyses concerning alertness before task performance, sleep quality and duration during the protocol, and vigilance or sleepiness during experimental SD night, see section 2 in supplementary material.

## 4. Discussion

In the present study, we used diffusion weighted imaging (DWI) to explore the dynamic microstructural reorganization happening in the short-term following learning and its long-term modulation by the presence or absence of sleep during the first post-training night. Also, we investigated the microstructural changes within minutes of practice on a previously experienced material.

Behaviourally, motor performance improved during learning and reached the same level for both groups at the end of Day 1. A similar pattern was observed during relearning at Day 5 both for the sleep deprived (SD) and regular sleep (RS) group. At morning retest at Day 2, RTs were slower in the SD than the RS group, which was predictable after a full night spent awake. Besides mere motor learning, RT significantly increased on random as compared to sequential blocks, indicating that both groups learned the repeated sequence adequately and similarly, that this knowledge persisted 3 days later, and that delayed practice at relearning on previously acquired material led to further performance improvement gains accumulating when re-practicing the task. No post-training sleep effect was detected. The evolution of performance with time and practice is consistent with previous findings [74], [75]. Concerning the absence of a sleep effect on delayed performance, other reports also found that post-learning sleep did not systematically result in increased performance at delayed retest [76]– [78]. Especially when focussing on the SRTT literature, sleep-related effects were not consistently observed. For example, differentiated neural patterns were detected using fMRI between post-learning sleep and sleep deprivation conditions 3 days after the experimental night, but in the absence of observable behavioural differences [79]. Similarly, other experimental protocols failed to detect sleep-specific effects on the optimization of sequential knowledge [80]–[82].

At the structural level, we found rapid learning-related modifications in brain’s microstructure after 1h of motor sequence training (vs. baseline), in line with previous reports showing changes even after more restricted learning times [30], [31]. In the cortical ribbon, changes were observed in a large part of the occipitoparietal and temporal area, respectively known for subtending visuomotor and somatosensory transformation and integration processes [83], [84]. Decreased MD was found in extended clusters in these areas, potentially reflecting increased tissue density. Also, decreased FWF in similar but more restricted territories suggests increased tissue proportion. Various cellular processes activated during learning may be responsible for increased tissue proportion and density. For instance, animal studies found gliogenesis as one of the key mechanisms activated during post-learning neocortical remodelling [85], [86]. Synaptogenesis and neuronal morphology changes in cortical areas such as the motor cortex have also been found [25], [87]–[89]. To a lesser extent, neurogenesis might also be involved although neocortical neurogenesis remains disputed nowadays [85], [90], [91]. However, increased NDI in regions exhibiting MD and FWF changes indicates that at least part of this reorganized/created cortical tissues may be neurites. Finally, ODI changes in a few clusters, again in analogous regions, also suggest tissue reorganization in the brain tissue microstructure, even if at a smaller magnitude. Interestingly, while MD decreases have been repeatedly reported in the neocortex following learning [28], [30], [92], a previous study using NODDI to investigate spatial learning-related structural plasticity observed FWF decreases but no NDI increase contrary to our results [92]. Besides task differences, this might be due to the fact we used the revised version [65], [66] instead of the original NODDI model [32] used in this preceding experiment, probably giving us the opportunity to gain in preciseness when it comes to disentangling the different components influencing MD alterations. Then, in subcortical regions of interest, decreased MD was found in the left putamen, right thalamus, and caudate, bilateral hippocampus and cerebellar cortex, together with increased FA in the right caudate and cerebellar cortex. Additionally, analyses on NODDI parameters highlighted reduced FWF in bilateral thalamus, hippocampus, and cerebellar cortex, similarly to MD changes, enhanced NDI in left putamen, and hippocampus and increased ODI in left hippocampus. MD reductions in the hippocampus and cerebellum after motor practice confirm previously reported motor learning-related microstructural changes developing in the short term [30], [31]. Also, sprouting of new mossy fibre terminals was observed following learning in mice hippocampus [93]. Increased NDI in the hippocampus and other ROIs is in line with these findings, whereas modifications in FA suggest increased directionality, and changes in ODI rapid motor learning-related reorganization. Altogether, modifications in structural parameters suggest a rapid motor learning-related remodelling in most of our ROIs and a set of neocortical regions encompassing a large part of the occipitoparietal and temporal cortices, reflecting learning-related neuronal brain plasticity.

Then, we hypothesized that delayed post-learning changes would be modulated by the presence/absence of post-learning sleep, and that this would be reflected in the first DWI acquisition on Day 5 compared to baseline. Small but significant clusters persisted in the cortical ribbon, but no learning-related changes were found in ROIs in the long term. Also, sleep did not modulate these changes: both groups exhibited the same pattern at Day 5 (DWI_3_) than Day 1 (DWI_1_). As discussed above, it is possible that consolidation for a sequential motor SRTT (or at least this one) relies more on time than sleep, both at the behavioural and structural level. Alternatively, it cannot be excluded that microstructural modifications take more time than a few days to be fully consolidated and observable, or that SD participants benefitted from the recovery nights after the RS/SD experimental night to catch up with the sleep group before the delayed scan, as such effect was already reported at the behavioural level [94]. Diffusion MRI scans obtained at the outset of the RS/SD night experimental night in the morning of Day 2 were not suitable for investigating this issue due to circadian confounds [47], [95] and contamination by the lack of sleep [96]–[98].

Lastly, we hypothesized that microstructural parameters would change again when re-exposed to the initial motor task, and that these changes could be modulated by the sleep opportunity on the post-learning night, as shown a in a prior study investigating topographical learning [92]. In the cortical ribbon, the pattern was similar to the one observed during the first learning episode but with a smaller magnitude, and this was not affected by the sleep manipulation. At the functional level, cortical involvement was shown to increase with sequential proficiency, whereas hippocampus and dorsomedial striatum progressively disengage [12]. In the case of the relearning session at Day 5, the structural network was already partially shaped during the first learning episode at Day 1, and thus only needed to be refined and fine-tuned during re-exposure, which likely explains the smaller amplitude of the modifications. Finally, correlations between DWI metrics and behavioural measures were all non-significant, at variance with previous reports disclosing correlations between behaviour and WM [20] or GM [20], [27] modifications. However, correlations in those past studies were found exclusively in the hippocampus for GM [20], [27] and in the fornix for WM [20] after 2h of spatial training. Both areas are heavily solicited and recruited when performing spatial tasks. In our case, a more widespread network seemed to be recruited to answer the motor task demands, maybe explaining the fact no strong correlation emerged between one specific region and motor performance.

Due to heavy demands entailed by this protocol and the fact that post-learning sleep deprivation was our experimental manipulation parameter, we chose not to include a motor control task. At the end of the protocol, participants practiced random and sequential blocks in the fMRI setting to partially disentangle the brain areas subtending motor and sequential components of performance in the SRTT. Functional MRI results evidenced higher sequence-related activation in the bilateral precuneus, caudate and left hippocampus, and a stronger involvement of the visuo-motor and executive areas during random practice. Interestingly, the precuneus also exhibited structural modifications in MD, NDI, FWF and ODI following the learning and relearning session as well as persisting NDI and ODI changes at day 5 in the right precuneus only. The right caudate showed MD decrease and FA increase after learning and the hippocampus showed bilateral MD and FWF decrease, right FA increase and left NDI increase following learning right NDI increase following relearning. These results confirm not only the repeatedly observed overlap between structurally and functionally engaged regions [18], [19], [28], suggesting that momentary adaptations in functional connectivity alters structural connections, which in turn affect functional connectivity [18], but they also corroborate the fact our results are sequence-specific at least in the precuneus, caudate and hippocampus, rather than attributable to motor practice only.

To sum up, we observed important and rapid tissue remodelling in response to sequential motor learning and relearning in occipitoparietal and temporal regions, as well as in ROIs involved in motor processing. However, we found limited persistence of those changes 3 days after initial learning. The use of the revised NODDI model in combination with conventional DTI brought us one step closer to image the cellular mechanisms and more specifically, the axonal and dendritic remodelling present in humans in response to learning independently from the glial adaptation that happens simultaneously to answer the task demands. However, no sleep effect was detected 3 days later, suggesting that either sleep-related cellular changes are too subtle to be identified at that macroscopic level using non-invasive DWI measures, or that consolidation in the SRTT we used is not sleep-dependent. These issues should be investigated in further studies. Nonetheless, the use of DTI combined with NODDI, or other complex biophysical models opens the way to reunite the cellular processes underlying learning- and sleep-related remodelling observed only in animals until now, with non-invasive brain imaging techniques applicable to humans.

## 5. Acknowledgments and Funding

W.S is supported by the Fonds de la Recherche Scientifique (F.R.S.-F.N.R.S., Aspirant Research Fellowship). The study and the postdoctoral fellows A.L. and M.G. were also supported by the F.N.R.S. and the Fonds Wetenschappelijk Onderzoek – Vlaanderen (F.W.O.) under the Excellence of Science (EOS) Project (MEMODYN, No. 30446199 to P.P. and H.Z.). The authors thank Brecht Somers, Julie Galinier, and Katja Skoda for their help in data acquisition.

## 6. Author contributions

Conceptualization, W.S. and P.P.; methodology, W.S., P.P. and T.V.; software, H.Z. and W.S.; formal analysis, W.S., A.L. and M.G.; investigation, W.S.; resources, P.P.; writing—original draft preparation, W.S.; writing—review and editing, P.P., H.Z., A.L., T.V. and M.G.; visualization, W.S. and A.L.; supervision, P.P. and H.Z.; funding acquisition, P.P. All authors have read and agreed to the published version of the manuscript.

