## Supplementary Material for "Microstructural dynamics of motor learning and sleep-dependent consolidation: a diffusion imaging study"

#### 1. Detailed Behavioural Results

For motor learning at Day 1, the ANOVA computed on mean reaction time (RT) for sequential blocks performed with within-subject factor *Block* (1:25, 27:30) and between-subject factor *Sleep* (RS vs. SD) disclosed a main *Block* effect ($F_{4.584, 270.445} = 109.153$, $p < 0.001$, $\eta_p^2 = 0.649$) characterized by progressive decrease in mean RT with task practice. Also, as expected, no main *Sleep* ($F_{1, 59} = 0.121$, $p = 0.729$, $\eta_p^2 = 0.002$) nor *Block*Sleep* interaction ($F_{4.584, 270.445} = 0.414$, $p = 0.823$, $\eta_p^2 = 0.007$) was found as the experimental manipulation did not happen yet at this stage. Thus, both groups exhibited a similar RT decrease over task practice in the initial learning session. Additionally, a separate ANOVA was conducted with within-subject factor *Block* type (sequential blocks 25 & 27 vs random block 26) and between-subject factor *Sleep* (RS vs. SD). This analysis disclosed a main *Block* effect ($F_{1.791, 105.691} = 227.806$, $p < 0.001$, $\eta_p^2 = 0.794$), and post-hoc tests showed that RT in pseudo-random block 26 was significantly slower than in sequential block 25 ($p < 0.001$) and 27 ($p < 0.001$). It indicates that participants learned the sequence and started anticipating the upcoming position in sequential blocks, and that performance improvement was not merely due to motor practice. As previously, no main effect of *Sleep* ($F_{1, 59}$ = 0.069, $p$ = 0.793, $\eta_p^2$ = 0.001) or *Block*Sleep* interaction ($F_{1.791, 105.691}$ = 0.002, $p$ = 0.997, $\eta_p^2$ = 3.032e$^{-5}$) was found. Similar ANOVAs were performed on accuracy measures. No main *Block* ($F_{14.509, 856.019}$ = 1.102; $p$ = 0.351, $\eta_p^2$ = 0.018), *Sleep* ($F_{1, 59}$ = 2.350, $p$ = 0.131, $\eta_p^2$ = 0.038), or *Block*Sleep* interaction ($F_{14.509, 856.019}$ = 1.143, $p$ = 0.314, $\eta_p^2$ = 0.019) effects were found across sequential blocks, indicating that accuracy remained stable over the learning session. Also, no difference in accuracy was found when comparing pseudo-random block 26 with sequential blocks 25 or 27 (main *Block* effect: $F_{2, 118}$ = 2.521, $p$ = 0.085, $\eta_p^2$ = 0.041; main *Sleep* effect $F_{1, 59}$ = 0.462, $p$ = 0.499, $\eta_p^2$ = 0.008; *Block*Sleep* interaction ($F_{2, 118}$ = 3.022, $p$ = 0.052, $\eta_p^2$ = 0.049).

For morning retest at Day 2, the mixed ANOVA looking at the evolution between the mean RT of the 2 last blocks of the learning session (LS D1; blocks 29:30) and the 2 blocks performed during retest on day 2 after the experimental night (RE D2; blocks 31:32) between-subject factor Sleep (RS vs. SD) disclosed no main *Day* ($F_{1, 59}$ = 0.154, $p$ = 0.696, $\eta_p^2$ = 0.003) or *Group* effect ($F_{1, 59}$ = 0.872, $p$ = 0.354, $\eta_p^2$ = 0.015). The *Day*Sleep* interaction was significant ($F_{1, 59}$ = 7.840, $p$ = 0.007, $\eta_p^2$ = 0.117) but post-hoc analyses did not reveal any significant comparison after Bonferroni correction (all $p$s > 0.173). Concerning accuracy, a similar ANOVA revealed no *Sleep* ($F_{1, 59}$ = 0.043, $p$ = 0.836, $\eta_p^2$ = 7.328e$^{-4}$), nor *Day*Sleep* interaction ($F_{1, 59}$ = 1.334, $p$ = 0.253, $\eta_p^2$ = 0.022). However, there was a main *Day* effect ($F_{1, 59}$ = 12.995, $p$ < 0.001, $\eta_p^2$ = 0.180) with a significantly lower accuracy at the beginning of the retest at D2 compared to the end of the learning session at Day 1.

For delayed motor memory consolidation at Day 5, the ANOVA compared mean performance on the 2 last blocks of the learning session (LS D1; blocks 29:30) and the 2 first blocks of the relearning session (RL D5; blocks 33:34) with within-subject factor *Day* (End LS D1 vs. Begin RL D5) and between-subject factor *Sleep* (RS vs. SD) to assess delayed offline gains in performance. The analysis revealed a significant decrease in mean RT between D1 and D5 ($F_{1, 59}$ = 13.968, $p$ < 0.001, $\eta_p^2$ = 0.191). However, both groups exhibited a similar decrease over time with no significant *Sleep* ($F_{1, 59}$ = 0.125, $p$ = 0.725, $\eta_p^2$ = 0.002) nor *Day*Sleep* interaction ($F_{1, 59}$ = 0.317, $p$ = 0.575, $\eta_p^2$ = 0.005) effect. Also, no significant effect was found concerning changes in accuracy (all $p$s > 0.343).

Looking at motor sequence relearning at Day 5, an ANOVA on mean reaction time (RT) for all sequential blocks with within-subject factor *Block* (33:34, 36:47, 49:52; random blocks being numbered 35 & 48) and between-subject factor *Sleep* (RS vs. SD) disclosed a main *Block* effect ($F_{9.441, 557.011} = 28.864$, $p < 0.001$, $\eta_p^2 = 0.329$) with a decrease in mean RT over the sequential blocks. However, no main *Sleep* ($F_{1, 59} = 0.784$, $p = 0.380$, $\eta_p^2 = 0.013$), nor *Block*Sleep* interaction ($F_{9.441, 557.011} = 1.106$, $p = 0.356$, $\eta_p^2 = 0.018$) was found, suggesting that post-learning sleep availability did not impact the behavioural time course for the practice on previously learned material. Besides, two separate ANOVAs comparing both pseudo-random blocks with their preceding and following sequential block respectively were conducted with within-subject factor *Block* (34:36 or 47:49) and between-subject factor *Sleep* (RS vs. SD). Both analyses disclosed slower RTs in pseudo-random blocks 35 and 48 than in the surrounding sequential blocks (all *p*s < 0.001). Regarding accuracy measures, no main *Block* ($F_{10.379, 612.382} = 1.112$; $p = 0.349$, $\eta_p^2 = 0.019$), *Sleep* ($F_{1, 59} = 2.219$, $p = 0.142$, $\eta_p^2 = 0.036$), or *Block*Sleep* interaction ($F_{10.379, 612.382} = 1.078$, $p = 0.377$, $\eta_p^2 = 0.018$) effects were found across sequential blocks. Also, no difference in accuracy was found when comparing pseudo-random block 35 with sequential blocks 34 or 36 (all *p*s > 0.828), neither when comparing pseudo-random block 48 with sequential blocks 47 or 49 (all *p*s > 0.132).

Also, we computed a complementary mixed ANOVA comparing sequential-specific amelioration at the end of learning and at the beginning of relearning (respectively computed as the difference in RT between the mean of the 2 sequential blocks surrounding random block 26 or 35 minus the random block, then divided by the random block and multiplied by 100) with between-subject factor *Sleep* (RS vs. SD). The analysis revealed a significant difference ($F_{1, 59} = 54.097$, $p < 0.001$, $\eta_p^2 = 0.478$) with a significantly larger difference between random and sequential bloc at the beginning of RL (-46,653%) compared to the end of LS (-37.219%), confirming the previous findings showing effective delayed memory consolidation. However, no main *Sleep* ($F_{1, 59} = 0.237$, $p = 0.628$, $\eta_p^2 = 0.004$) or *Sleep*Day* interaction ($F_{1, 59} = 3.488$, $p = 0.067$, $\eta_p^2 = 0.056$) effects were found.

Finally, all participants reported noticing the sequential nature of the task by the end of the procedure.

### 2. Additional (control) Analyses

#### 2.1. Alertness before task performance

An ANOVA computed on Reciprocal Reaction Time (RRT) in the PVT5 (1/RT [1] with within-subject factor *PVT Session* (D1, D2, D5) and between-subject factor *Sleep* (SD vs. RS) found no main *Sleep* effect ($F_{1, 59} = 2.492$, $p = 0.120$, $\eta_p^2 = 0.041$), but a significant main *PVT Session* ($F_{1.632, 96.299} = 14.351$, $p < 0.001$, $\eta_p^2 = 0.196$) and a *Sleep*PVT Session* interaction ($F_{1.632, 96.299} = 24.053$, $p < 0.001$, $\eta_p^2 = 0.290$) effects. As expected, post hoc tests disclosed similar alertness in both groups before the learning session at Day 1 ($p = 1.000$) or the relearning session at Day 5 ($p = 1.000$), and a strong difference ($p < 0.001$) at morning retest at Day 2 at the outset of the experimental night, with significantly lower alertness in the SD than in the RS group.

#### 2.2. Sleep Quality and Duration during the protocol

The ANOVA performed with within-subject factor *Night Quality* (1:3, 5:7; night 4 being the experimental night) and between-subject factor *Sleep* (SD vs. RS) disclosed a main effect of *Night Quality* ($F_{5, 290} = 34.055$, $p < 0.001$, $\eta_p^2 = 0.370$) and *Night Quality*Sleep* interaction ($F_{5, 290} = 3.314$, $p = 0.006$, $\eta_p^2 = 0.054$) with a significantly lower sleep quality and this even more for the RS group, on the first night corresponding to our habituation night at the lab, but a significantly higher sleep quality on the 5$^{th}$ night, thus the night following the experimental night spend at the lab. However, no main effect of *Sleep* ($F1_{, 58} = 2.916$, $p = 0.093$, $\eta_p^2 = 0.048$) was highlighted.

A similar ANOVA conducted on sleep duration with within-subject factor *Night* (1:3, 5:7) and between-subject factor *Sleep* (SD vs. RS) revealed no main *Sleep* effect ($F_{1, 58} = 0.332$, $p = 0.567$, $\eta_p^2 = 0.006$) nor *Night*Sleep* interaction ($F_{3.986, 231.196} = 1.176$, $p = 0.322$, $\eta_p^2 = 0.020$). There was a main *Night* effect $F_{3.986, 231.196} = 8.832$, $p < 0.001$, $\eta_p^2 = 0.132$) with reduced sleep duration on the 5$^{th}$ night (i.e., the night after the experimental RS/SD night, *ps* < 0.001).

#### 2.3. Sleep Quality and Duration on the experimental RS night

Separate ANOVAs were carried out in the RS group only to verify if the quantity/quality of the experimental night was equivalent to the previous and next night. The ANOVA on sleep quality with within-subject factor *Night* (3:5) revealed a main *Night* ($F_{2, 58} = 11.700$, $p < 0.001$, $\eta_p^2 = 0.287$) effect with a lower quality on the experimental night compared to the previous ($p = 0.005$) and next ($p < 0.001$) ones, that can be explained by sleeping with the hd-EEG setting on that experimental night. The ANOVA on sleep duration disclosed no significant difference between the 3 nights ($F_{2, 58} = 2.813$, $p = 0.068$, $\eta_p^2 = 0.088$).

#### 2.4. Vigilance and Sleepiness during experimental SD night

The ANOVA performed on sleepiness KSS-scores with within-subject factor *Time* (hourly score between 22h and 8h) evidenced a *Time* effect ($F_{3.686, 106.883} = 24.080$, $p < 0.001$, $\eta_p^2 = 0.454$) with increasing sleepiness all over the night. Similarly, the ANOVA computed on vigilance RRT ($PVT_{10}$) scores with within-subject factor Time (bi-hourly score between 22h and 6h) evidenced a Time effect ($F_{1.585, 45.977} = 16.668$, $p < 0.001$, $\eta_p^2 = 0.365$) with decreasing alertness over the night.

### Table 1. Participant's demographics

| | Sleep Deprivation (SD) Group<br>n = 31 | Regular Sleep (RS) Group<br>n = 30 | Total sample<br>n = 61 |
|---|---|---|---|
| Pittsburgh Sleep Quality Index (sleep quality) | mean score = 3.32<br>SD = 1.14<br>min = 1<br>max = 6 | mean score = 3.33<br>SD = 1.35<br>min = 0<br>max = 5 | mean score = 3.33<br>SD = 1.24<br>min = 0<br>max = 6 |
| Morningness–Eveningness Questionnaire (chronotype) | mean score = 54.71<br>SD = 8.17<br>min = 37<br>max = 73 | mean score = 53.42 SD = 7.56<br>min = 32<br>max = 68 | mean score = 54.07<br>SD = 7.84<br>min = 32<br>max = 73 |
| Edinburgh Inventory (laterality) | mean score = 5.68<br>SD = 6.16<br>min = -10<br>max = 10 | mean score = 4.27<br>SD = 6.64<br>min = -10<br>max = 10 | mean score = 4.98<br>SD = 6.39<br>min = −10<br>max = 10 |
| Gender | ♀ = 16<br>♂ = 15 | ♀ = 15<br>♂ = 15 | ♀ = 31<br>♂ = 30 |
| Age | mean score = 21.07<br>SD = 2.35<br>min = 19<br>max = 29 | mean score = 21.57<br>SD = 2.16<br>min = 18<br>max = 29 | mean age = 21.31<br>SD = 2.26<br>min = 18<br>max = 29 |

*Table 1.* Detailed information concerning sleep quality, chronotype, laterality, gender, and age with mean (mean score), standard deviation (SD), minimum (min) and maximum (max) for every group and in the total sample.

### Table 2. Learning-related short-term structural changes (Day 1; DWI1 vs. DWI2)in the cortical ribbon

| Annot | Size ($mm^2$) | WghtVtx | NVtxs | nbClust |
|---|---|---|---|---|
| MD – right hemisphere | | | | |
| paracentral | 1876.55 | -20159.60 | 4473.00 | 2.00 |
| precentral | 616.85 | -5965.65 | 1468.00 | 4.00 |
| lateraloccipital | 544.64 | -2719.04 | 728.00 | 4.00 |
| precuneus | 479.97 | -5693.94 | 1207.00 | 7.00 |
| parsopercularis | 436.19 | -3919.29 | 953.00 | 3.00 |
| inferiorparietal | 369.79 | -2966.48 | 710.00 | 6.00 |
| postcentral | 366.52 | -3676.02 | 888.00 | 5.00 |
| supramarginal | 310.32 | -3113.13 | 729.00 | 5.00 |
| middletemporal | 295.63 | -2810.67 | 690.00 | 3.00 |
| insula | 259.82 | -2359.87 | 606.00 | 4.00 |
| superiorparietal | 253.84 | -2241.31 | 544.00 | 3.00 |
| rostralanteriorcingulate | 236.20 | -1682.28 | 439.00 | 1.00 |
| fusiform | 224.53 | -1603.51 | 374.00 | 4.00 |
| transversetemporal | 212.70 | -3391.37 | 663.00 | 1.00 |
| superiorfrontal | 189.67 | -1429.12 | 359.00 | 5.00 |
| lingual | 156.16 | -735.18 | 200.00 | 6.00 |
| cuneus | 74.33 | -289.16 | 79.00 | 2.00 |
| bankssts | 41.64 | -536.90 | 146.00 | 1.00 |
| isthmuscingulate | 38.31 | -389.62 | 109.00 | 2.00 |
| parahippocampal | 33.36 | -206.37 | 58.00 | 1.00 |
| rostralmiddlefrontal | 21.38 | -116.40 | 37.00 | 3.00 |
| superiortemporal | 12.97 | -94.28 | 27.00 | 1.00 |
| inferiortemporal | 10.50 | -58.79 | 19.00 | 3.00 |
| pericalcarine | 7.88 | -25.42 | 8.00 | 1.00 |
| parsorbitalis | 4.51 | -15.40 | 5.00 | 1.00 |
| MD – left hemisphere | | | | |
| inferiorparietal | 3113.02 | -24785.42 | 5441.00 | 4.00 |
| precuneus | 1556.79 | -16318.41 | 3474.00 | 4.00 |
| insula | 1227.16 | -13460.19 | 3301.00 | 2.00 |
| precentral | 1008.95 | -10708.41 | 2562.00 | 4.00 |
| lingual | 890.26 | -5684.58 | 1309.00 | 4.00 |
| superiorparietal | 611.86 | -6586.74 | 1615.00 | 6.00 |
| superiorfrontal | 510.69 | -4231.93 | 1092.00 | 7.00 |
| caudalanteriorcingulate | 327.71 | -2775.57 | 717.00 | 1.00 |
| superiortemporal | 271.84 | -2492.47 | 671.00 | 3.00 |
| caudalmiddlefrontal | 270.79 | -2613.18 | 540.00 | 2.00 |
| inferiortemporal | 263.16 | -1854.19 | 453.00 | 2.00 |
| cuneus | 219.79 | -1143.72 | 315.00 | 2.00 |
| posteriorcingulate | 218.10 | -2400.21 | 611.00 | 2.00 |
| supramarginal | 198.77 | -1610.50 | 461.00 | 8.00 |
| postcentral | 190.90 | -1782.27 | 450.00 | 4.00 |

| | | | | |
|---|---|---|---|---|
| fusiform | 163.05 | -1054.55 | 293.00 | 3.00 |
| transversetemporal | 148.39 | -1209.77 | 338.00 | 1.00 |
| lateralorbitofrontal | 134.69 | -808.33 | 224.00 | 2.00 |
| paracentral | 78.00 | -712.31 | 209.00 | 4.00 |
| rostralanteriorcingulate | 73.47 | -522.23 | 152.00 | 3.00 |
| bankssts | 59.67 | -364.20 | 110.00 | 2.00 |
| lateraloccipital | 52.21 | -211.00 | 67.00 | 4.00 |
| rostralmiddlefrontal | 41.77 | -280.14 | 76.00 | 2.00 |
| middletemporal | 30.59 | -242.79 | 65.00 | 1.00 |
| isthmuscingulate | 14.99 | -127.06 | 37.00 | 2.00 |
| parahippocampal | 12.46 | -90.54 | 27.00 | 1.00 |
| NDI – right hemisphere | | | | |
| precuneus | 775.46 | 8043.37 | 1929.00 | 7.00 |
| lateraloccipital | 629.95 | 3463.61 | 872.00 | 7.00 |
| superiorfrontal | 389.26 | 3628.63 | 901.00 | 5.00 |
| inferiorparietal | 371.53 | 2583.22 | 660.00 | 10.00 |
| superiortemporal | 367.83 | 3140.96 | 824.00 | 6.00 |
| precentral | 333.47 | 3817.95 | 931.00 | 5.00 |
| middletemporal | 288.47 | 2404.63 | 656.00 | 7.00 |
| fusiform | 258.61 | 1717.82 | 457.00 | 5.00 |
| rostralanteriorcingulate | 192.08 | 1286.13 | 341.00 | 2.00 |
| parsopercularis | 188.86 | 1687.68 | 459.00 | 1.00 |
| supramarginal | 145.85 | 1617.02 | 405.00 | 7.00 |
| insula | 140.17 | 1241.41 | 314.00 | 2.00 |
| parsorbitalis | 138.58 | 708.70 | 183.00 | 1.00 |
| caudalanteriorcingulate | 120.41 | 1134.42 | 290.00 | 1.00 |
| medialorbitofrontal | 113.68 | 525.47 | 159.00 | 2.00 |
| lateralorbitofrontal | 70.74 | 475.24 | 134.00 | 2.00 |
| lingual | 49.36 | 202.04 | 61.00 | 3.00 |
| rostralmiddlefrontal | 44.98 | 311.98 | 91.00 | 2.00 |
| parahippocampal | 32.81 | 199.39 | 61.00 | 1.00 |
| caudalmiddlefrontal | 27.54 | 204.29 | 62.00 | 2.00 |
| inferiortemporal | 25.81 | 105.86 | 32.00 | 1.00 |
| postcentral | 22.30 | 247.56 | 71.00 | 1.00 |
| paracentral | 22.05 | 203.57 | 63.00 | 2.00 |
| pericalcarine | 22.03 | 120.52 | 35.00 | 1.00 |
| parstriangularis | 17.99 | 98.50 | 31.00 | 1.00 |
| transversetemporal | 9.48 | 97.24 | 31.00 | 1.00 |
| superiorparietal | 6.34 | 40.24 | 13.00 | 1.00 |
| posteriorcingulate | 4.45 | 30.76 | 10.00 | 1.00 |
| NDI – left hemisphere | | | | |
| lateraloccipital | 1544.80 | 9355.97 | 2347.00 | 6.00 |
| posteriorcingulate | 1057.23 | 10645.27 | 2485.00 | 2.00 |
| caudalmiddlefrontal | 1050.05 | 10022.21 | 2208.00 | 1.00 |
| inferiorparietal | 1034.10 | 8625.82 | 2006.00 | 8.00 |
| lingual | 912.92 | 6601.82 | 1483.00 | 3.00 |

| | | | | |
|---|---|---|---|---|
| supramarginal | 691.96 | 6492.45 | 1704.00 | 9.00 |
| superiorfrontal | 480.57 | 3552.27 | 941.00 | 10.00 |
| precentral | 438.50 | 3907.95 | 973.00 | 5.00 |
| rostralmiddlefrontal | 400.34 | 2644.02 | 695.00 | 6.00 |
| insula | 367.67 | 4354.00 | 1051.00 | 3.00 |
| postcentral | 314.11 | 3053.48 | 825.00 | 4.00 |
| fusiform | 259.21 | 1797.07 | 456.00 | 1.00 |
| middletemporal | 249.88 | 2037.41 | 554.00 | 4.00 |
| superiortemporal | 178.28 | 1404.50 | 388.00 | 4.00 |
| bankssts | 162.96 | 1517.72 | 349.00 | 1.00 |
| lateralorbitofrontal | 157.98 | 988.03 | 253.00 | 5.00 |
| precuneus | 115.93 | 836.87 | 239.00 | 5.00 |
| caudalanteriorcingulate | 114.02 | 872.23 | 242.00 | 2.00 |
| superiorparietal | 105.60 | 699.51 | 204.00 | 6.00 |
| parsopercularis | 88.64 | 584.26 | 157.00 | 1.00 |
| medialorbitofrontal | 60.28 | 235.33 | 65.00 | 1.00 |
| rostralanteriorcingulate | 46.13 | 320.69 | 92.00 | 2.00 |
| parstriangularis | 38.68 | 205.09 | 63.00 | 3.00 |
| parsorbitalis | 15.26 | 71.85 | 22.00 | 1.00 |
| transversetemporal | 7.57 | 67.31 | 21.00 | 1.00 |
| cuneus | 4.90 | 22.12 | 7.00 | 1.00 |
| FWF – right hemisphere | | | | |
| superiorfrontal | 179.87 | -1333.69 | 352.00 | 1.00 |
| superiorparietal | 136.38 | -942.39 | 253.00 | 3.00 |
| fusiform | 79.25 | -465.31 | 126.00 | 2.00 |
| superiortemporal | 73.21 | -645.99 | 165.00 | 2.00 |
| caudalmiddlefrontal | 71.27 | -437.81 | 122.00 | 2.00 |
| lateraloccipital | 65.78 | -289.36 | 85.00 | 3.00 |
| postcentral | 64.43 | -312.30 | 92.00 | 1.00 |
| paracentral | 54.60 | -438.39 | 130.00 | 2.00 |
| lingual | 53.37 | -318.62 | 93.00 | 3.00 |
| precuneus | 48.89 | -274.58 | 88.00 | 5.00 |
| transversetemporal | 36.58 | -366.28 | 94.00 | 1.00 |
| insula | 35.57 | -244.77 | 72.00 | 2.00 |
| cuneus | 32.14 | -90.31 | 29.00 | 2.00 |
| medialorbitofrontal | 18.00 | -129.16 | 41.00 | 1.00 |
| caudalanteriorcingulate | 16.98 | -122.30 | 35.00 | 1.00 |
| middletemporal | 15.99 | -85.34 | 26.00 | 1.00 |
| precentral | 15.28 | -108.46 | 33.00 | 2.00 |
| posteriorcingulate | 8.70 | -73.27 | 22.00 | 2.00 |
| FWF – left hemisphere | | | | |
| lateraloccipital | 405.75 | -2277.28 | 602.00 | 6.00 |
| lingual | 226.43 | -1336.56 | 341.00 | 1.00 |
| precentral | 197.19 | -1884.72 | 534.00 | 4.00 |
| superiorparietal | 192.67 | -1356.88 | 386.00 | 5.00 |
| cuneus | 130.60 | -693.99 | 191.00 | 2.00 |

| | | | | |
|---|---|---|---|---|
| postcentral | 104.04 | -762.21 | 225.00 | 6.00 |
| insula | 82.03 | -952.39 | 251.00 | 2.00 |
| precuneus | 59.87 | -443.42 | 120.00 | 3.00 |
| caudalanteriorcingulate | 59.85 | -530.53 | 142.00 | 2.00 |
| superiortemporal | 56.48 | -477.35 | 136.00 | 3.00 |
| inferiorparietal | 49.23 | -373.70 | 105.00 | 4.00 |
| fusiform | 39.44 | -211.27 | 64.00 | 2.00 |
| superiorfrontal | 29.01 | -256.15 | 79.00 | 1.00 |
| pericalcarine | 20.12 | -94.02 | 30.00 | 2.00 |
| supramarginal | 18.86 | -128.88 | 41.00 | 3.00 |
| isthmuscingulate | 11.93 | -102.73 | 29.00 | 1.00 |
| rostralmiddlefrontal | 6.99 | -43.72 | 13.00 | 1.00 |
| paracentral | 4.33 | -36.98 | 12.00 | 2.00 |
| ODI – right hemisphere | | | | |
| caudalmiddlefrontal | 43.56 | 178.07 | 54.00 | 1.00 |
| lateralorbitofrontal | 36.35 | -195.07 | 53.00 | 1.00 |
| inferiorparietal | 23.90 | -64.56 | 62.00 | 2.00 |
| precuneus | 23.04 | 158.20 | 46.00 | 1.00 |
| superiortemporal | 16.06 | -100.43 | 29.00 | 1.00 |
| superiorfrontal | 13.95 | -111.17 | 34.00 | 2.00 |
| postcentral | 11.64 | 104.82 | 31.00 | 1.00 |
| fusiform | 8.44 | -54.57 | 17.00 | 1.00 |
| supramarginal | 8.37 | 73.27 | 22.00 | 1.00 |
| middletemporal | 2.67 | 16.07 | 5.00 | 1.00 |
| ODI – left hemisphere | | | | |
| postcentral | 94.41 | 546.67 | 232.00 | 4.00 |
| posteriorcingulate | 17.68 | -157.69 | 48.00 | 1.00 |
| precuneus | 16.18 | 121.89 | 35.00 | 1.00 |
| precentral | 11.38 | -93.06 | 29.00 | 2.00 |
| rostralmiddlefrontal | 7.55 | 38.01 | 12.00 | 1.00 |
| caudalmiddlefrontal | 7.47 | -6.02 | 12.00 | 2.00 |
| superiorfrontal | 6.74 | 34.83 | 11.00 | 1.00 |
| lateraloccipital | 2.51 | 9.07 | 3.00 | 1.00 |
| superiortemporal | 1.57 | -9.06 | 3.00 | 1.00 |

*Table 2.* Clusters showing significant MD, NDI, FWF and ODI changes in both groups following learning at Day 1 after cluster-wise correction gathered by cortical region. Annot = the name of the regions the maximum cluster value falls into using Desikan-Killiany atlas. Size ($mm^2$) = sum of the cluster sizes in millimeters square. WghtVtx = sum of the cluster weights (size*intensity), a negative weight indicate decrease in intensity while a positive value points towards an increase in intensity. NVtxs = sum of the number of vertices in the clusters from the region. NbClust = number of clusters in the labelled area.

**Table 3. Delayed structural changes and sleep-related effects in Pre-learning vs. Pre-relearning (DWI1 vs. DWI3 x RS vs. SD) in the cortical ribbon**

| Annot | Size ($mm^2$) | WghtVtx | NVtxs | nbClust |
|---|---|---|---|---|
| MD – right hemisphere | | | | |
| parstriangularis | 24.59 | 160.02 | 47.00 | 1.00 |
| superiorparietal | 20.35 | 232.05 | 62.00 | 1.00 |
| bankssts | 4.19 | 24.61 | 8.00 | 1.00 |
| MD – left hemisphere | | | | |
| lingual | 19.17 | -48.31 | 34.00 | 2.00 |
| superiortemporal | 8.56 | -31.68 | 10.00 | 1.00 |
| NDI – right hemisphere | | | | |
| superiorparietal | 30.67 | -250.41 | 93.00 | 2.00 |
| superiortemporal | 25.27 | -197.15 | 54.00 | 1.00 |
| lingual | 13.94 | -60.88 | 19.00 | 2.00 |
| parsorbitalis | 13.14 | 50.33 | 16.00 | 1.00 |
| fusiform | 9.29 | -53.67 | 16.00 | 1.00 |
| paracentral | 4.93 | 30.66 | 10.00 | 1.00 |
| precuneus | 4.54 | 37.65 | 12.00 | 1.00 |
| parsopercularis | 1.94 | 9.18 | 3.00 | 1.00 |
| lateralorbitofrontal | 1.71 | -9.17 | 3.00 | 1.00 |
| NDI – left hemisphere | | | | |
| superiortemporal | 37.97 | 202.21 | 60.00 | 3.00 |
| supramarginal | 18.86 | 152.52 | 47.00 | 1.00 |
| rostralmiddlefrontal | 6.51 | 30.94 | 10.00 | 1.00 |
| postcentral | 6.31 | 52.95 | 17.00 | 2.00 |
| lateralorbitofrontal | 1.53 | 9.16 | 3.00 | 1.00 |
| FWF – right hemisphere | | | | |
| superiorparietal | 28.05 | -145.11 | 44.00 | 2.00 |
| lateraloccipital | 12.40 | 60.87 | 19.00 | 1.00 |
| superiorparietal | 6.66 | -54.50 | 17.00 | 1.00 |
| precentral | 5.89 | 47.60 | 15.00 | 1.00 |
| supramarginal | 4.15 | -34.95 | 11.00 | 1.00 |
| fusiform | 2.92 | -18.37 | 6.00 | 1.00 |
| FWF – left hemisphere | | | | |
| lingual | 25.91 | -140.26 | 41.00 | 1.00 |
| superiortemporal | 23.34 | -110.41 | 36.00 | 3.00 |
| caudalmiddlefrontal | 11.88 | 61.15 | 19.00 | 2.00 |
| fusiform | 5.15 | -24.78 | 8.00 | 1.00 |
| ODI – right hemisphere | | | | |
| rostralmiddlefrontal | 27.53 | -126.29 | 39.00 | 2.00 |
| superiorfrontal | 25.81 | -123.56 | 35.00 | 1.00 |
| parahippocampal | 17.25 | -114.21 | 34.00 | 1.00 |
| inferiorparietal | 13.09 | 30.96 | 31.00 | 2.00 |
| precuneus | 9.34 | -70.91 | 22.00 | 1.00 |

| | | | | |
|---|---|---|---|---|
| inferiortemporal | 8.31 | -55.37 | 17.00 | 1.00 |
| postcentral | 1.41 | 15.35 | 5.00 | 1.00 |
| ODI – left hemisphere | | | | |
| lateraloccipital | 38.26 | 196.80 | 58.00 | 2.00 |
| superiorfrontal | 31.38 | 159.01 | 46.00 | 2.00 |
| postcentral | 19.34 | -146.35 | 43.00 | 1.00 |
| lateralorbitofrontal | 15.44 | -113.09 | 33.00 | 1.00 |
| inferiorparietal | 2.55 | -12.06 | 4.00 | 1.00 |
| fusiform | 2.51 | -15.28 | 5.00 | 1.00 |
| precentral | 1.69 | 9.03 | 3.00 | 1.00 |
| supramarginal | 1.25 | -12.20 | 4.00 | 1.00 |

*Table 3.* Clusters exhibiting significant MD, NDI, FWF and ODI modifications at Day 5 when compared to baseline after cluster-wise correction brought together by region. The table represent changes for both groups as no group difference was found. Annot = the name of the regions the maximum cluster value falls into using Desikan-Killiany atlas. Size ($mm^2$) = sum of the cluster sizes in millimeters square. WghtVtx = sum of the cluster weights (size*intensity), a negative weight indicate decrease in intensity while a positive value points towards an increase in intensity. NVtxs = sum of the number of vertices in the clusters from the region. NbClust = number of clusters in the labelled area.

**Table 4. Post-relearning structural changes and sleep-related effects (Day 5; DWI3 vs. DWI4 X RS vs. SD) in the cortical ribbon.**

| Annot | Size ($mm^2$) | WghtVtx | NVtxs | nbClust |
|---|---|---|---|---|
| MD – right hemisphere | | | | |
| bankssts | 389.58 | -4585.77 | 1113.00 | 1.00 |
| lateraloccipital | 386.21 | -2399.85 | 569.00 | 4.00 |
| insula | 362.85 | -4391.45 | 1077.00 | 4.00 |
| superiorparietal | 279.84 | -3128.42 | 695.00 | 3.00 |
| cuneus | 219.73 | -933.86 | 271.00 | 4.00 |
| precuneus | 160.36 | -1583.66 | 444.00 | 3.00 |
| posteriorcingulate | 157.52 | -1678.58 | 429.00 | 3.00 |
| supramarginal | 108.67 | -1222.79 | 337.00 | 2.00 |
| lingual | 97.17 | -364.38 | 106.00 | 1.00 |
| lateralorbitofrontal | 80.31 | -660.23 | 176.00 | 3.00 |
| medialorbitofrontal | 77.69 | -392.65 | 99.00 | 1.00 |
| precentral | 77.14 | -784.93 | 204.00 | 3.00 |
| superiorfrontal | 71.21 | -627.06 | 189.00 | 3.00 |
| fusiform | 65.53 | -398.57 | 112.00 | 2.00 |
| postcentral | 55.29 | -575.90 | 162.00 | 4.00 |
| middletemporal | 34.97 | -292.38 | 87.00 | 1.00 |
| paracentral | 33.69 | -337.72 | 97.00 | 3.00 |
| caudalmiddlefrontal | 31.15 | -216.74 | 66.00 | 4.00 |
| inferiorparietal | 25.48 | -160.06 | 51.00 | 1.00 |
| isthmuscingulate | 23.36 | -216.88 | 59.00 | 1.00 |
| superiortemporal | 9.69 | -63.51 | 20.00 | 1.00 |
| parahippocampal | 8.72 | -40.40 | 13.00 | 2.00 |
| rostralmiddlefrontal | 2.37 | -15.23 | 5.00 | 1.00 |
| MD – left hemisphere | | | | |
| precuneus | 751.53 | -7370.88 | 1646.00 | 5.00 |
| insula | 348.72 | -3763.13 | 977.00 | 4.00 |
| superiorparietal | 311.65 | -2336.47 | 605.00 | 5.00 |
| inferiorparietal | 294.56 | -1699.66 | 486.00 | 4.00 |
| cuneus | 144.87 | -636.28 | 195.00 | 3.00 |
| lateraloccipital | 143.42 | -782.17 | 218.00 | 3.00 |
| postcentral | 137.59 | -1322.00 | 378.00 | 8.00 |
| superiorfrontal | 118.33 | -886.17 | 231.00 | 4.00 |
| supramarginal | 94.01 | -943.42 | 272.00 | 2.00 |
| caudalmiddlefrontal | 91.89 | -691.05 | 201.00 | 2.00 |
| precentral | 78.48 | -801.16 | 228.00 | 2.00 |
| superiortemporal | 73.07 | -735.21 | 198.00 | 3.00 |
| lingual | 55.36 | -247.83 | 69.00 | 2.00 |
| posteriorcingulate | 50.08 | -448.39 | 138.00 | 3.00 |
| rostralmiddlefrontal | 48.75 | -287.72 | 83.00 | 2.00 |
| lateralorbitofrontal | 39.16 | -275.47 | 76.00 | 2.00 |
| bankssts | 37.05 | -355.27 | 100.00 | 1.00 |

| | | | | |
|---|---|---|---|---|
| pericalcarine | 36.96 | -337.14 | 95.00 | 1.00 |
| transversetemporal | 36.38 | -463.49 | 103.00 | 1.00 |
| fusiform | 16.72 | -148.96 | 41.00 | 1.00 |
| isthmuscingulate | 1.52 | -24.80 | 8.00 | 1.00 |
| NDI – right hemisphere | | | | |
| lateraloccipital | 388.86 | 2360.88 | 569.00 | 5.00 |
| caudalmiddlefrontal | 311.03 | 2307.11 | 664.00 | 5.00 |
| medialorbitofrontal | 214.50 | 1259.92 | 305.00 | 4.00 |
| superiorparietal | 163.18 | 1655.70 | 429.00 | 2.00 |
| fusiform | 149.64 | 1037.97 | 296.00 | 2.00 |
| insula | 144.12 | 1678.32 | 454.00 | 2.00 |
| posteriorcingulate | 112.20 | 1049.94 | 294.00 | 3.00 |
| lateralorbitofrontal | 103.73 | 722.46 | 210.00 | 3.00 |
| bankssts | 101.05 | 1223.97 | 319.00 | 2.00 |
| rostralmiddlefrontal | 74.06 | 535.02 | 147.00 | 1.00 |
| cuneus | 58.65 | 239.56 | 71.00 | 1.00 |
| precentral | 56.59 | 408.83 | 125.00 | 4.00 |
| superiortemporal | 51.28 | 387.86 | 107.00 | 2.00 |
| middletemporal | 50.96 | 325.43 | 92.00 | 3.00 |
| precuneus | 45.70 | 435.08 | 131.00 | 3.00 |
| isthmuscingulate | 28.66 | 231.21 | 66.00 | 1.00 |
| superiorfrontal | 26.07 | 253.37 | 73.00 | 1.00 |
| inferiorparietal | 23.99 | 179.20 | 52.00 | 2.00 |
| paracentral | 14.50 | 133.84 | 40.00 | 1.00 |
| parsopercularis | 2.80 | 18.32 | 6.00 | 1.00 |
| NDI – left hemisphere | | | | |
| precuneus | 769.05 | 7579.75 | 1783.00 | 1.00 |
| rostralmiddlefrontal | 282.92 | 1836.29 | 495.00 | 3.00 |
| superiorfrontal | 181.55 | 1267.05 | 360.00 | 4.00 |
| insula | 158.21 | 1738.50 | 426.00 | 3.00 |
| superiortemporal | 127.55 | 1124.10 | 311.00 | 4.00 |
| lateralorbitofrontal | 114.75 | 860.87 | 238.00 | 4.00 |
| parahippocampal | 108.50 | 842.85 | 235.00 | 1.00 |
| precentral | 97.35 | 889.18 | 252.00 | 2.00 |
| postcentral | 68.58 | 712.04 | 187.00 | 3.00 |
| inferiorparietal | 54.74 | 432.62 | 120.00 | 4.00 |
| bankssts | 52.53 | 455.58 | 131.00 | 3.00 |
| superiorparietal | 47.77 | 450.90 | 127.00 | 3.00 |
| lateraloccipital | 44.37 | 183.12 | 58.00 | 3.00 |
| supramarginal | 32.71 | 319.36 | 94.00 | 2.00 |
| parsopercularis | 26.41 | 202.34 | 65.00 | 1.00 |
| caudalmiddlefrontal | 19.05 | 102.00 | 33.00 | 2.00 |
| entorhinal | 17.06 | 125.30 | 37.00 | 1.00 |
| middletemporal | 16.79 | 93.03 | 28.00 | 1.00 |
| paracentral | 8.02 | 81.10 | 25.00 | 1.00 |
| posteriorcingulate | 6.87 | 54.09 | 17.00 | 1.00 |

| | | | | |
|---|---|---|---|---|
| inferiortemporal | 5.86 | 34.22 | 11.00 | 1.00 |
| FWF – right hemisphere | | | | |
| cuneus | 151.30 | -542.60 | 163.00 | 3.00 |
| lateraloccipital | 55.55 | -345.21 | 84.00 | 2.00 |
| superiorparietal | 48.03 | -503.17 | 131.00 | 3.00 |
| postcentral | 33.28 | -370.22 | 110.00 | 2.00 |
| precentral | 22.65 | -237.66 | 73.00 | 4.00 |
| insula | 11.49 | -127.98 | 36.00 | 1.00 |
| precuneus | 9.16 | -90.92 | 28.00 | 1.00 |
| middletemporal | 7.85 | -50.48 | 16.00 | 1.00 |
| paracentral | 6.42 | -44.46 | 14.00 | 1.00 |
| superiorfrontal | 6.17 | -55.63 | 17.00 | 1.00 |
| supramarginal | 4.52 | -33.99 | 11.00 | 1.00 |
| superiortemporal | 2.66 | -22.55 | 7.00 | 1.00 |
| bankssts | 1.74 | -15.35 | 5.00 | 1.00 |
| caudalanteriorcingulate | 1.49 | -9.10 | 3.00 | 1.00 |
| FWF – left hemisphere | | | | |
| lateraloccipital | 101.74 | -627.88 | 160.00 | 2.00 |
| pericalcarine | 78.85 | -656.81 | 170.00 | 3.00 |
| cuneus | 73.55 | -271.82 | 83.00 | 2.00 |
| lingual | 68.98 | -292.88 | 78.00 | 1.00 |
| precuneus | 58.09 | -368.62 | 106.00 | 3.00 |
| postcentral | 54.88 | -543.09 | 151.00 | 3.00 |
| precentral | 45.20 | -511.50 | 140.00 | 2.00 |
| superiorfrontal | 38.11 | -197.89 | 54.00 | 1.00 |
| superiorparietal | 30.13 | -226.53 | 68.00 | 3.00 |
| insula | 25.79 | -248.79 | 75.00 | 3.00 |
| inferiorparietal | 22.77 | -152.88 | 45.00 | 2.00 |
| rostralmiddlefrontal | 16.10 | 58.42 | 18.00 | 1.00 |
| medialorbitofrontal | 8.94 | 43.25 | 14.00 | 1.00 |
| lateralorbitofrontal | 6.61 | 58.47 | 19.00 | 1.00 |
| posteriorcingulate | 3.79 | -28.07 | 9.00 | 1.00 |
| ODI – right hemisphere | | | | |
| precuneus | 53.28 | 364.10 | 107.00 | 4.00 |
| postcentral | 51.51 | 481.72 | 131.00 | 3.00 |
| lateraloccipital | 35.53 | 124.15 | 38.00 | 1.00 |
| superiorfrontal | 32.23 | 83.85 | 63.00 | 3.00 |
| precentral | 30.54 | 219.43 | 63.00 | 2.00 |
| inferiorparietal | 15.96 | 63.37 | 38.00 | 3.00 |
| superiorparietal | 15.07 | 81.35 | 36.00 | 3.00 |
| cuneus | 8.01 | 37.92 | 12.00 | 1.00 |
| insula | 4.94 | 38.82 | 12.00 | 1.00 |
| paracentral | 4.43 | -36.14 | 11.00 | 1.00 |
| rostralmiddlefrontal | 1.08 | -6.06 | 2.00 | 1.00 |
| middletemporal | 0.53 | 3.01 | 1.00 | 1.00 |
| ODI – left hemisphere | | | | |

| | | | | |
|---|---|---|---|---|
| postcentral | 42.35 | 387.63 | 109.00 | 3.00 |
| superiorfrontal | 25.00 | 192.59 | 57.00 | 2.00 |
| lateraloccipital | 13.06 | -61.30 | 19.00 | 1.00 |
| precuneus | 9.81 | 65.34 | 21.00 | 2.00 |
| entorhinal | 4.14 | 30.69 | 10.00 | 1.00 |

*Table 4.* Clusters indicating significant MD, NDI, FWF and ODI alterations following relearning at Day 5 after cluster-wise correction compiled by region. The table represent changes for both groups as no group difference was found. Annot = the name of the regions the maximum cluster value falls into using Desikan-Killiany atlas. Size ($mm^2$) = sum of the cluster sizes in millimeters square. WghtVtx = sum of the cluster weights (size*intensity), a negative weight indicate decrease in intensity while a positive value points towards an increase in intensity. NVtxs = sum of the number of vertices in the clusters from the region. NbClust = number of clusters in the labelled area.
